# Aerobic exercise reverses aging-induced depth-dependent decline in cerebral microcirculation

**DOI:** 10.1101/2023.02.12.528244

**Authors:** Paul Shin, Qi Pian, Hidehiro Ishikawa, Gen Hamanaka, Emiri T Mandeville, Guo Shuzhen, Fu Buyin, Mohammed Alfadhel, Srinivasa Rao Allu, Ikbal Şencan-Eğilmez, Baoqiang Li, Chongzhao Ran, Sergei A Vinogradov, Cenk Ayata, Eng H Lo, Ken Arai, Anna Devor, Sava Sakadžić

## Abstract

Aging is a major risk factor for cognitive impairment. Aerobic exercise benefits brain function and may promote cognitive health in older adults. However, underlying biological mechanisms across cerebral gray and white matter are poorly understood. Selective vulnerability of the white matter to small vessel disease and a link between white matter health and cognitive function suggests a potential role for responses in deep cerebral microcirculation. Here, we tested whether aerobic exercise modulates cerebral microcirculatory changes induced by aging. To this end, we carried out a comprehensive quantitative examination of changes in cerebral microvascular physiology in cortical gray and subcortical white matter in mice (3-6 vs. 19-21 months old), and asked whether and how exercise may rescue age-induced deficits. In the sedentary group, aging caused a more severe decline in cerebral microvascular perfusion and oxygenation in deep (infragranular) cortical layers and subcortical white matter compared with superficial (supragranular) cortical layers. Five months of voluntary aerobic exercise partly renormalized microvascular perfusion and oxygenation in aged mice in a depth-dependent manner, and brought these spatial distributions closer to those of young adult sedentary mice. These microcirculatory effects were accompanied by an improvement in cognitive function. Our work demonstrates the selective vulnerability of the deep cortex and subcortical white matter to aging-induced decline in microcirculation, as well as the responsiveness of these regions to aerobic exercise.

## 1. Introduction

The cerebral white matter is significantly affected by aging.(Cees De Groot et al., 2000; de Leeuw et al., 2001) Previous work suggests that, in comparison with superficial gray matter, subcortical white matter has greater tissue loss due to aging and is more susceptible to hypoperfusion.(Gunning-Dixon et al., 2009; Hase et al., 2019; Li et al., 2020) Indeed, white matter lesions might impair cognitive function more than gray matter lesions,(Reber et al., 2021) underscoring the importance of understanding the biological mechanisms of aging related white matter degeneration for targeted interventions.

Cerebral small vessel disease (CSVD) refers to a range of pathological processes affecting small arterioles, capillaries, and venules supplying the white matter and deep structures of the brain. CSVD is a common cause of stroke and an important contributor to age-related cognitive decline and dementia.(Pantoni, 2010; Smith & Markus, 2020) Most CSVD related strokes affect subcortical white and deep gray matter. In addition, there is increased awareness of the role of CSVD in accelerating the pathogenesis of Alzheimer’s disease (AD), with some studies suggesting the possibility that white matter changes are the starting point of AD.(Defrancesco et al., 2014; Esiri et al., 1999; Radanovic et al., 2013; Snowdon et al., 2011) Despite this growing awareness, our mechanistic understanding of aging-related changes in microcirculation as a function of depth in cortex and underlying white matter is incomplete.

Aerobic exercise is a promising strategy to improve neurocognitive function in aging and reduce the risk of age-related neurological disorders.(D. E. Barnes et al., 2003; Yaffe et al., 2001) Most studies have thus far focused on exercise-responsive molecules that could lead to improvement of neural physiology and, thereby, cognitive performance.(de Miguel et al., 2021; Islam et al., 2021; Valenzuela et al., 2020; Wang & Holsinger, 2018) Yet, how exercise may exert beneficial effects on vascular contributions to brain aging remain to be fully understood. In particular, it is unclear how exercise normalizes cerebral microcirculation, and whether there are differential effect across brain regions, including the gray and white matter.

In this study, we compared young (3-6 month old) vs. old (19-21 months old) mice to assess the effects of normal aging, and then asked whether 5 months of voluntary aerobic exercise can alter or rescue brain microcirculation in old 20-month-old mice. Cerebral microvascular perfusion and oxygenation in the cortex and subcortical white matter was quantified with two-photon microscopy (2PM) and optical coherence tomography (OCT) in awake mice.

Our results showed age-related declines in capillary red-blood-cell (RBC) flux and capillary oxygen partial pressure (pO_2_) in the deep cortical layers and subcortical white matter, while voluntary exercise improved these measures, including the cerebral blood flow (CBF) in cortical ascending venules. Interestingly, the regions that experienced the highest decline were also the ones that benefited the most from the exercise. These microcirculatory effects were accompanied by an improvement in cognitive function. Our results may provide insights into how sedentary aging and aerobic exercise affect cerebral microvascular physiology and particularly emphasize the physiologic importance of effects in the deep cortex and subcortical white matter.

## 2. Methods

### 2.1 Animal preparation and experimental protocol

All animal surgical and experimental procedures were conducted following the Guide for the Care and Use of Laboratory Animals and approved by the Massachusetts General Hospital Subcommittee on Research Animal Care. All efforts were made to minimize the number of animals used and their suffering, in accordance with the Animal Research: Reporting in Vivo Experiments (ARRIVE) guidelines. Female C57BL/6N mice (n=18, 12 months old) were obtained from National Institute on Aging colonies. The experimental timeline of the study is shown in Figure 1a. A chronic cranial window (round shape, 3 mm in diameter) was implanted on the left hemisphere, centered over the E1 whisker barrel (2.0 mm posterior to bregma and 3.0 mm lateral from the midline) at the age of 13 months. Mice were allowed four weeks to recover fully from surgery. Next, mice were randomly divided into two groups (exercise: n=9; sedentary: n=9), and all mice were housed individually. Cages of mice in the exercise group were equipped with wireless running wheels (ENV-047; Med Associates) that could be used by mice at any time. The total number of wheel rotations was recorded daily through a wireless hub device (DIG-807; Med Associates). For each mouse in the exercise group, the wheel running activity was characterized based on the intensity of exercise, defined as the average daily running distance over five months (Figures 1b,c). After five months of voluntary exercise (or standard housing for the sedentary controls), mice underwent restraint training for awake imaging. Mice in the exercise group were continuously housed in cages with running wheels during both training and imaging weeks. All mice were gradually habituated to longer restraint periods, up to 2 hours. They were rewarded with sweetened milk every ∼15 min during training and imaging. Optical measurements were performed between 19 and 21 months of age. The measurements were performed during 5-6 weeks in the following order: two-photon phosphorescence pO_2_ imaging, 2PM angiography, optical intrinsic signal imaging (OISI), 2PM imaging of capillary RBC flux, and Doppler OCT. 7-10 days of a break was given between measurements in each animal. Finally, behavioral tests were performed, and blood samples were collected from the animals for hematocrit (Hct) measurements before the animals were euthanized.

**Figure 1.**
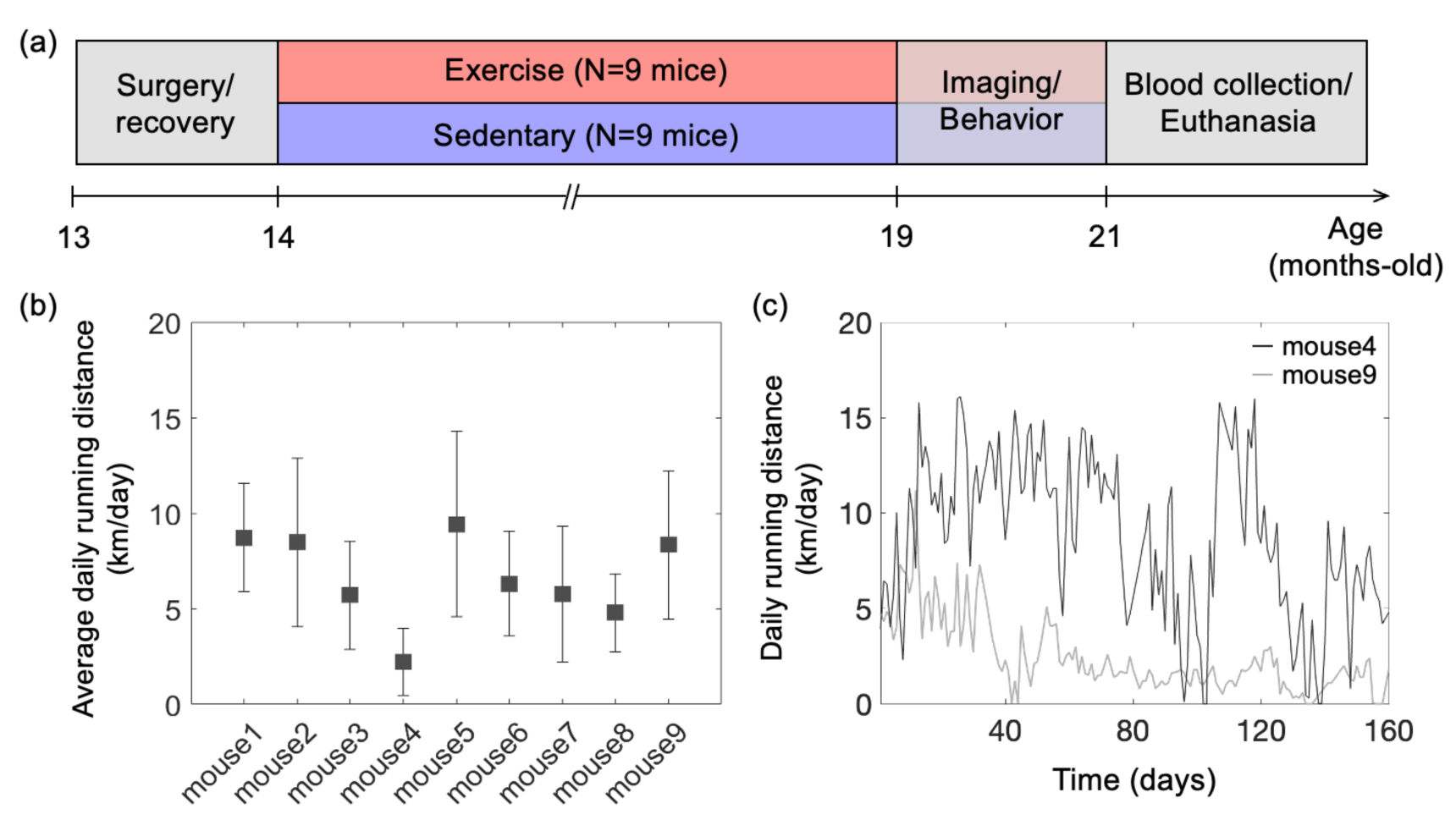
Animal preparation and experiment design. (a) Timeline of the study. Optical measurements and behavioral testing were performed after 5 months of voluntary exercise when the animals were 19-21 months of age. (b) Average daily running distance in km per day for each mouse in the exercise group, calculated as the sum of daily running distance divided by a total running period of 5 months. Data are shown as mean ± SD. (c) Daily running distance for two representative mice in the exercise group across time.

Some measurements in the old mice were compared with similar measurements in the young adult mice. Female C57BL/6N mice, 6-month-old (n=8) for OISI (OISI group) and 3-month-old (n=3) for capillary RBC flux imaging (2PM group), were used in this work. Chronic cranial window implantation, animal recovery from the surgery, and restraint training were performed following the same protocols as in the old mice. Therefore, measurements were performed at 7 months in the OISI group and 4 months in the 2PM group.

### 2.2 Multimodal optical imaging in awake mice at rest

In this study, we employed our previously described home-built multimodal imaging system that features multiple optical imaging capabilities, including 2PM, OCT, and OISI.^3^ Figure 2a shows a schematic of the multimodal imaging system. All optical measurements were made in the head-restrained awake mice at rest. The measurements were conducted while mice were resting on a suspended soft fabric bed in a home-built imaging platform. An accelerometer was attached to the suspended bed to monitor the signals induced by animal motion. Data affected by the motion were rejected during data processing based on the signal generated by the accelerometer. The data acquired when the accelerometer reading exceeded the threshold value, which was determined empirically by comparing signals obtained during stationary and movement periods, were rejected from the analysis. During the experiments, animal behavior was continuously monitored using a web camera (LifeCam Cinema; Microsoft) with a LED illumination at 940 nm, and a reward (sweetened milk) was offered every ∼15 min. Experiments were terminated if signs of discomfort or anxiety were observed.

**Figure 2.**
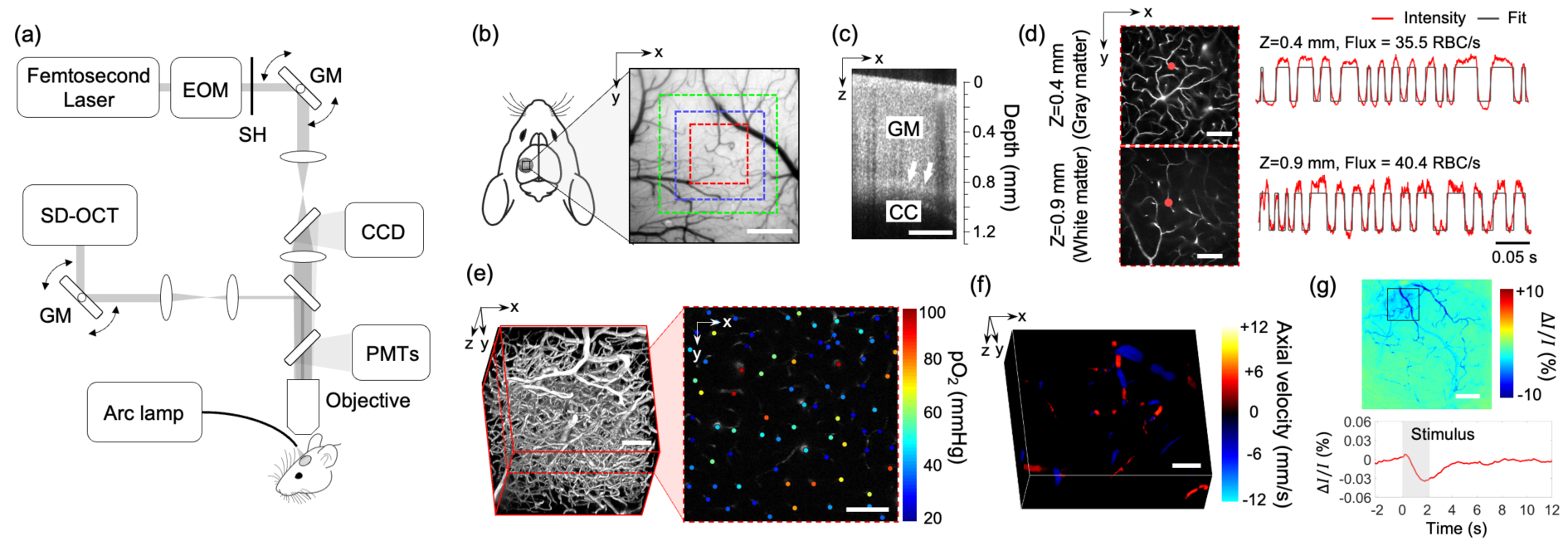
Experimental setup and imaging protocols. (a) Schematic of our home-built multimodal optical system featuring primary components of the system. A 50 kHz spectral-domain OCT system was designed to partially share the imaging optics with the two-photon microscope. A Hg:Xe arc lamp in combination with a CCD camera was used for OISI. EOM: electro-optic modulator, SH: shutter, GM: galvanometer mirror pair. (b) A CCD image of brain surface vasculature in the mouse barrel cortex showing the ROIs where various optical measurements were performed (red ROI: capillary RBC flux, pO_2_, and microvasculature imaging, blue ROI: Doppler OCT imaging, and green ROI: OCT intensity imaging). (c) A representative OCT intensity B-scan image extracted from a volumetric OCT image. White arrows indicate the boundary between the gray matter (GM) and the corpus callosum (CC), which appears as a bright band in the image. (d) Survey scan images of cerebral microvasculature of the region outlined with the red square in (b) obtained by two-photon microscope at two imaging depths (z=0.4 and 0.9 mm). Two representative fluorescent intensity time courses acquired within the capillaries at the locations indicated by the red dots in the survey angiograms are presented on the right. (e) A 3D angiogram of the mouse cortex acquired by the two-photon microscope at the location outlined by the red square in (b). One representative 2D plane from the angiogram acquired at a depth of 200 µm showing pO_2_ measurements from different capillary segments. pO_2_ values were color-coded (in mmHg) and spatially co-registered with the angiogram. (f) A 3D Doppler OCT image showing an axial velocity map of the diving vessels at the location outlined by the blue square in (b). (g) An OIS image of the cranial window obtained by calculating the relative intensity difference between the post-stimulus response image and pre-stimulus baseline. The region of activation is manually selected from the OIS image as indicated by a black square. The lower panel shows a time course of the relative intensity change due to sensory-evoked hemodynamic response induced by a 2-s-long whisker stimulation, averaged over the selected region of interest. Scale bars: 400 µm for (b) and (c), 100 µm for (d), (e), and (f), and 500 µm for (g).

### 2.3 Two-photon microscope for fluorescence and phosphorescence imaging

A two-photon microscope integrated into our multimodal imaging system was used for capillary RBC flux measurements and 3D microvascular angiography(Li et al., n.d.; Sakadžić et al., 2010). A pulsed laser (InSight DeepSee; Spectra-Physics, tuning range: 680 nm to 1300 nm, ∼120 fs pulse width, 80 MHz pulse repetition rate) was used as an excitation light source. Laser power was controlled by an electro-optic modulator (EOM) (350-160; ConOptics, Inc.). The laser beam was focused by a water immersion objective lens (XLUMPLFLN20XW; Olympus) and scanned in the transverse (X-Y) plane by a pair of galvanometer mirrors (Saturn 5B; Pangolin Laser System, Inc.). The objective lens was translated along the Z-axis by a motorized stage (M-112.1EG; Physik Instrumente) to probe different depths ranging from the brain surface to depths beyond 1 mm below the surface. Emitted fluorescence was directed to detectors by an epi-dichroic mirror (FF875-Di01-38.1×51; Semrock Inc.) positioned above the objective, followed by an infrared blocker (FF01-890/SP-50; Semrock Inc.). The four-channel detector consists of four photomultiplier tubes (PMTs) paired with emission filters that cover a wide range of emission wavelengths. One detector channel with a PMT exhibiting a red-shifted spectral response (H10770PA-50; Hamamatsu) and a 709/167 nm emission filter was connected to a discriminator (C9744; Hamamatsu) and used for capillary RBC flux measurements. Another channel with a multialkali photocathode PMT (R3896; Hamamatsu) and a 525/50 nm emission filter was used for two-photon angiography.

We employed another home-built two-photon microscope for intravascular pO_2_ measurements. The second 2PM employs a Ti:Sapphire mode-locked laser (Mai Tai HP; Spectral Physics, tuning range: 690 nm to 1040 nm, ∼100 fs pulse width, 80 MHz pulse repetition rate) as a light source. The output laser power delivered to the sample is modulated with EOM (350-105-02; ConOptics Inc.). The transverse scanning was achieved by a two-axis 7-mm galvanometer scanner (6210H; Cambridge Technology). The laser beam was relayed to the back focal plane of the objective through the combination of a scan lens (f = 51 mm, 2x AC508-100-B, Thorlabs) and a tube lens (f =180 mm, Olympus). The Z-axis movement of the objective was controlled by two translation stages (ZFM2020 and PFM450E; Thorlabs). The phosphorescence emission signal from the sample was filtered by a dichroic mirror (FF875-Di01-38.1X51; Semrock Inc.) and an emission filter (FF01-795/150-25; Semrock Inc.) before detected by a PMT (H10770PA-50; Hamamatsu) and a photon counting unit (C9744; Hamamatsu).

The objective lens was heated by an electric heater (TC-HLS-05; Bioscience Tools) throughout the measurement to maintain the temperature of the water between the objective lens and the cranial window at 36-37 ℃.

### 2.4 Spectral-domain OCT

A low-coherence superluminescent diode with a center wavelength of 1300 nm was used as the light source (S5FC-1018S; Thorlabs). The line scan camera was operated with a 50 kHz acquisition rate (GL2048L; Sensors Unlimited). The sample arm of the system partially shares the imaging optics with the two-photon microscope, but it utilizes separate scanning optics, as shown in Figure 2a. The system has an axial resolution of 10 µm in tissue. The transverse resolution is 7 µm using a 10× objective lens (Mitutoyo Plan Apo NIR; Edmund Optics). The incident optical power on the sample was 5 mW. For OCT Doppler measurements, the maximum measurable flow speed without phase wrapping was ±12 mm/s when the direction of flow was parallel to the OCT beam and the minimum detectable velocity was ±0.7 mm/s which was determined by the phase noise of the system, measured as ±0.2 radians.

### 2.5 OISI system

A Hg:Xe arc lamp (66883; Newport) was used in combination with a band-pass filter (570/10 nm) for OISI. A 4× objective lens (XLFLUOR4X/340; Olympus) was used to achieve a wide field-of-view (FOV) that covered the entire cranial window. Two-dimensional images of the cranial window were acquired using a CCD camera with 100 ms exposure time (acA1300; Basler).

### 2.6 Measurements of capillary RBC flux, speed, and line-density in gray and white matter

Before imaging, the dextran-conjugated Alexa680 solution (70 kDa, 0.1-0.15 ml at 5% W/V in PBS; Thermo Fisher Scientific) was retro-orbitally injected into the bloodstream. Figure 2b shows a CCD image of the mouse cranial window positioned on the left somatosensory cortex. A volumetric OCT scan was performed with a FOV of 1×1 mm^2^ over the region shown with a green square box in Figure 2b. The acquired OCT volume was used to identify the boundary between the gray and white matter and measure the cortical thickness as we previously described (Figure 2c).(Li et al., 2020) After confirming localization of the white matter, 2PM was performed over the same region as for the OCT imaging, but with a smaller FOV of 500 × 500 µm^2^ indicated by a red square region (Figure 2b). The RBC flux measurements in the gray matter were performed at cortical depths of 150 and 400 µm, which correspond to layers II/III and IV. In the white matter, measurements were performed at a depth of 0.9-1.1 mm. Awake imaging lasted less than 2 hours and the flux measurements were performed only at three depths due to the time spent performing the OCT imaging prior to the flux measurements. At each depth, a survey image of the vasculature was acquired by raster scanning the beam across the FOV (Figure 2d). Then we manually selected measurement locations inside all the capillary segments identified within the survey image. Capillaries were identified based on their network structure and, in this work, all branching vessels from diving arterioles and surfacing venules were defined as capillaries. The laser beam was parked at each location for 0.9 s and the fluorescence signal was detected in the photon-counting mode. The photon counts were binned into 300-µs-wide bins, resulting in a fluorescence intensity time course with 3,000 time points. Figure 2d shows representative fluorescence signal transients taken from two measurement locations at different depths. Following the procedures described in our previous study,(Li et al., 2020) the fluorescence signal time course was segmented with a binary thresholding approach (red curve in Figure 2d) and RBC flux was calculated by counting the number of valleys in the segmented curve normalized by the acquisition time. Following our previously described procedures,(Li et al., n.d.) RBC speed for each RBC-passage event (valley) in the segmented curve was estimated as v = D⁄Δt, where D is RBC diameter, assumed to be 6 µm, and Δt is the width of the valley. For each capillary, the speed values estimated from each of the valleys in the segmented curve were averaged to obtain the mean RBC speed. Finally, RBC line-density was calculated as the ratio between combined time duration of all valleys to the total duration of the entire time course.

### 2.7 Intravascular *pO_2_*imaging and calculation of *SO_2_*and depth-dependent OEF

A phosphorescent oxygen-sensitive probe Oxyphor2P was diluted in saline and retro-orbitally injected before imaging (0.05 ml at ∼80 µM).(Esipova et al., 2019) The pO_2_ imaging was performed using 950 nm excitation wavelength and with a FOV of 500 ×500 µm^2^ at the same cortical region where the RBC flux measurements were performed (Figure 2b). The measurements were performed at cortical depths from the surface to 450 µm depth with 50 µm interval between depth locations. At each depth, two-dimensional raster scan of phosphorescence intensity was performed to acquire a survey image. Then, we manually selected the measurement locations inside all diving arterioles, surfacing venules, and capillary segments visually identified within the survey image. Next, the focus of excitation laser beam was parked at each selected segment to excite the Oxyphor2P with a 10-µs-long excitation gate. The resulting emitted phosphorescence was acquired during 290-µs-long collection time. Such 300-µs-long cycle was repeated 2,000 times to generate an average phosphorescence decay curve. The averaged curve was fitted to a single-exponential decay, and the lifetime was converted into absolute pO_2_ using a Stern-Volmer-like expression obtained from independent calibrations.(Esipova et al., 2019; Li et al., n.d., 2020; Şencan et al., 2022) The oxygen saturation of hemoglobin (SO_2_) and the depth-dependent oxygen extraction fraction (DOEF) were computed following our previously described procedures.(Li et al., n.d.) Briefly, the SO_2_ was calculated using the Hill equation based on the measured pO_2_ and the DOEF was calculated as (SO_2,#_ − SO_2,$_)⁄SO_2,#_, where SO_2,#_ and SO_2,$_ represent the mean SO_2_ in the diving arterioles and surfacing venules in a given cortical layer, respectively.

### 2.8 Characterization of morphological changes in cortical capillaries

FITC-dextran (Thermo Fisher Scientific, 70 kDa) was diluted in saline and injected retro-orbitally before imaging (, 0.05 ml at 5% W/V). A three-dimensional imaging of the cortical vasculature was performed with 500 × 500 µm^2^ FOV to cover the same region of interest (ROI) used for capillary flux and pO_2_ measurements (Figure 2e). The microvascular stack was generated by repeatedly acquiring images with the axial steps of 1 µm up to a depth of 400 µm. In each animal, a smaller ROI (200 × 200 µm^2^) was manually selected to cover the region containing mostly capillaries (e.g., capillary bed area). A 3D microvasculature corresponding to the selected ROI was segmented using a vessel segmentation algorithm, VIDA,(Tsai et al., 2009) and the number of capillary segments per volume (e.g., vessel segment density) and the average capillary segment length per volume (e.g., vessel segment length density) were calculated. The vessel segment was defined as the part of the vessel between two consecutive branching points.

### 2.9 Quantitative flow measurements using Doppler OCT

A total of 20 Doppler OCT volumes were continuously acquired with a 750 × 750 µm^2^ FOV at the region indicated by a blue square box in Figure 2b. 10 volumes exhibiting minimal motion artefacts were selected and averaged to generate a single Doppler OCT volume. Each Doppler OCT volume was comprised of 300 B-scans, where 3,000 A-scans were acquired per B-scan. The Doppler volume yields a three-dimensional map of the z-projection of RBC velocity (Figure 2f). For each depth slice in the volume, we measured flow in each surfacing venule over the FOV by estimating the integral of the velocity projection over the vessel cross section following the protocol previously described.(Srinivasan et al., 2011) We used 3D angiograms acquired by 2PM (see Method 2.7 for more details) to estimate the diameter of the vessels, used for cross-correlation analysis between the flow and vessel diameter. The measured flow values in each vascular segment were averaged within the depth range of 50-100 µm. Flow data measured in diving arterioles were excluded from analysis due to their much higher flow compared to venular flow that often causes excessive phase wrapping and signal fading.(Koch et al., 2009)

### 2.10 OISI imaging of hemodynamic response to functional activation

Two-dimensional CCD images were continuously acquired with a CCD camera for 18 seconds. Five seconds after the onset of image acquisition, a whisker stimulus was applied at 3 Hz for 2 s. A total of 20 stimulation trials were repeated with an inter-stimulus interval of 25 seconds. Among 20 trials, the data acquired during excessive animal motion (∼10% of the measurements) were excluded from analysis based on a threshold criterion applied to the accelerometer recordings. Bulk motion artifacts in the acquired CCD images caused by small transverse movement (< ∼50 µm) were compensated by 2D cross-correlation based motion correction algorithm.(Guizar-Sicairos et al., 2008) After motion compensation, images were averaged over trials before computing the fractional intensity difference between the response and baseline images (Figure 2g) as described in our previous work.(Şencan et al., 2022)

### 2.11 Behavioral tests

A novel object recognition test (NORT) was performed to evaluate whether exercise improves short-term memory in old mice. Briefly, mice were allowed to explore two identical objects in a testing arena for 5 minutes and returned to their home cage for 30 minutes. Mice were then moved back to the arena with one of the objects replaced with a novel object and allowed to explore the objects for 5 minutes. Behavioral performance was evaluated by estimating the discrimination index (DI). The DI was defined as exploration time of the novel object normalized by combined exploration time of both objects and was calculated during four different time intervals in each animal: during first one, two, three, and five minutes of the object exploration.

Y-maze test was used to measure the willingness of animals to explore new environments and spatial working memory. The testing occurred in a Y-shaped maze with three arms at a 120° angle from each other. Mice were placed at the center of the maze and allowed to freely explore the three arms for 10 minutes. The number of arm entries and the number of triads (consecutive entries into three different arms) were measured to calculate the percentage of spontaneous alteration, defined as the ratio of the number of alternating triads to total number of arm entries -2.

### 2.12 Hct measurements

Hct measurements were carried out by a blood gas analyzer (OPTI CCA-TS2; OPTI Medical Systems) using a B-type cassette. Mice were anesthetized with 2% isoflurane, and then approximately 250 µl of venous blood was collected from the inferior vena cava. The collected blood was aliquoted into two 100 µl capillary tubes. The Hct measurement was performed two times with each blood sample and the average of the two measurements was reported.

### 2.13 Statistical analysis

Statistical analysis was carried out using t-test or ANOVA (MATLAB, MathWorks Inc.). P value less than 0.05 was considered statistically significant. Details about the statistical analysis are provided in the figure legends and text, where relevant. Boxplots show the median value with a black line and the mean value with a plus symbol (+). Each box spans between the 25^th^ and the 75^th^ percentile of the data, defined as interquartile range (IQR). Whiskers of the boxplots extend from the lowest datum within 1.5 times the IQR of the lower quartile of the data to the highest datum within 1.5 times the IQR of the highest quartile of the data. For each parameter measured in this study, the acquired values were averaged to obtain the mean value for each mouse, and then the mean values were averaged over mice. Sample size (i.e., n=9 mice for aged sedentary and exercise group) were selected based on the assumption that the most demanding one is to detect 30% difference between the mean capillary pO_2_ values (coefficient of variance=0.3, power=0.8, significance=0.05).

## 3. Results

### 3.1 Exercise mitigates age-related decline of capillary RBC flux and induces capillary flow homogenization in the subcortical white matter

To examine the impact of aging on cortical and subcortical microcirculation, we examined the spatial distribution of capillary RBC flux across cortical and subcortical regions in aged mice and compared the result with measurements from awake C57BL/6N female mice of younger age (4 months old).

Capillary RBC flux measurements were conducted using 2PM imaging of RBC-induced shadows within blood plasma labeled by the Alexa-680, allowing the deep imaging into the subcortical white matter down to a depth of ∼1.1 mm.(Li et al., n.d.) The mean capillary RBC flux in aged sedentary mice was significantly lower than that in young sedentary mice in both gray and white matter (Figure 3a,b). Importantly, while in young sedentary mice, the mean white matter RBC flux tended to be slightly higher than the gray matter RBC flux, the relationship appeared reversed in aged sedentary mice, where mean RBC flux in the white matter strongly tended to be lower than RBC flux in the gray matter. This suggests that the microcirculation in the white matter is affected more than microcirculation in the gray matter by age-related changes.

**Figure 3.**
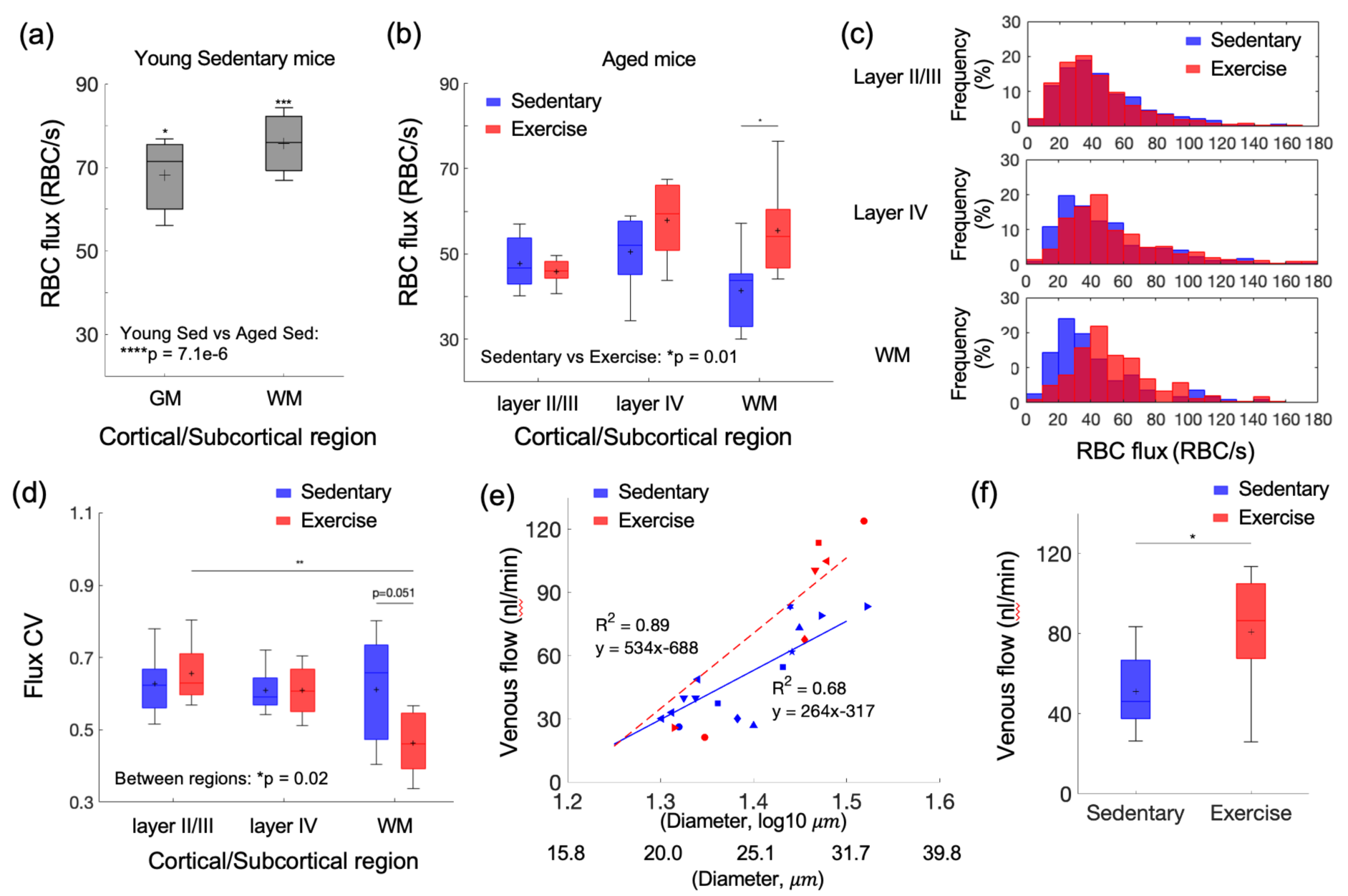
Aging and exercise induced alterations in cerebral microcirculation. (a) Mean capillary RBC flux in young-adult sedentary mice. The data are from 226 and 218 capillaries in three young sedentary mice in the gray matter (GM) and white matter (WM), respectively. Comparison was made between young and aged sedentary group (shown in (b)) in each gray (layers II/III, IV in aged sedentary group) and white matter region. (b) Capillary RBC flux across cortical layers II/III and IV, and subcortical white matter in aged sedentary and exercise group. (c) Histograms of capillary RBC flux in the gray and white matter in each animal group. (d) The coefficient of variance (CV) of capillary RBC flux across cortical layers II/III and IV, and subcortical white matter in sedentary and exercise group. (e) Venular flow versus vessel diameter. Different symbols represent different animals. The red dashed and blue solid line is the best fit result of each linear regression for sedentary and exercise group, respectively. (f) Mean venular flow in ascending venules in (d) in sedentary and exercise group. The data in (b-d) are from 921, 486, and 112 capillaries in 7 mice in the sedentary group and 1046, 465, and 238 capillaries in 8 mice in the exercise group, in cortical layers II/III, IV, and the white matter, respectively. The data in (e) and (f) are from 14 and 7 ascending venules in 9 and 6 mice in the sedentary and exercise group, respectively. Statistical analysis was carried out using Two-way ANOVA with post hoc Tukey’s in (a,b) and (d) and Student’s t-test in (f). *p<0.05; **p<0.01. Additional details on boxplots and animals excluded from the analyses are provided in the Supplementary document.

We next investigated whether voluntary exercise in aged mice exerts beneficial effects on capillary RBC flux across cortical layers and white matter. We found higher capillary flux in the exercise group compared with sedentary controls, which was most prominent in white matter (Figure 3b) and to a lesser extent in layer IV, while no change was observed in cortical layers II/III. Histograms of capillary RBC flux in gray and whiter matter confirmed this finding (Figure 3c). Exercise also tended to increase capillary RBC speed in layer IV of the cortex and the underlying white matter, while no change was detected in layers II/III (Figure 3 – Figure supplement 1). No significant difference in the RBC line-density between sedentary and exercise group was found (Figure 3 – Figure supplement 2). The RBC line-density of subcortical white matter was significantly higher than the RBC line-density of gray matter in both sedentary and exercise group, consistent with the previous numerical simulation result.(Gould et al., 2017)

Exercise decreased the coefficient of variation (CV) of capillary RBC flux in subcortical white matter but not the gray matter, suggesting more homogeneous microvascular blood flow in white matter in the exercise group (Figure 3d). In contrast, we did not find a difference in the CV of capillary RBC flux among the layers in the sedentary group.

Finally, we employed Doppler OCT to test the effect of exercise on the blood flow in larger vessels, particularly the ascending venules in the cortex. We performed a linear regression analysis between the venular blood flow and the logarithmic vessel diameter (Figure 3e). As expected, a strong correlation was found between the venular flow and the vessel diameter in both groups, consistent with the previous observation of positive correlation between venular flow speed and the vessel diameter in mice.(Santisakultarm et al., 2012) Importantly, the regression slope for the exercise group was steeper than that for the sedentary group (p=0.005, analysis of covariance, ANCOVA). Blood flow in ascending venules was significantly larger (by ∼ 46%) in the exercise group compared to sedentary controls. This result agreed with the improved capillary RBC flux in the white matter with exercise.

The capillary RBC flux in subcortical white matter was moderately negatively correlated with the average daily running distance (i.e., exercise intensity; R^2^ = 0.31). No strong correlation was found between other measured parameters and the exercise intensity (Figure 3 – Figure supplement 3).

### 3.2 Aging-induced reduced microvascular oxygenation in deeper cortical regions was rescued by 5-months of voluntary exercise

We next asked whether aging and exercise-induced changes in capillary RBC flux affected the microvascular oxygenation as well. We performed two-photon phosphorescence lifetime imaging using our two-photon microscope and a phosphorescent oxygen-sensitive probe (Oxyphor 2P) to examine variations of capillary mean pO_2_ in both sedentary and exercise group as a function of cortical depth (Figure 4a). We have previously shown that the resting-state capillary mean pO_2_ gradually increased from layer I to IV by ∼ 6 mmHg in young-adult mice (3-5 months old) of the same strain and sex.(Li et al., n.d.) In contrast, the aged sedentary mice here exhibited a different pattern, with the capillary mean pO_2_ reaching a plateau in layers II/III (capillary mean pO_2_, layer I: 42±1 mmHg, layers II/III: 46±1 mmHg, layer IV: 45±2 mmHg).

**Figure 4.**
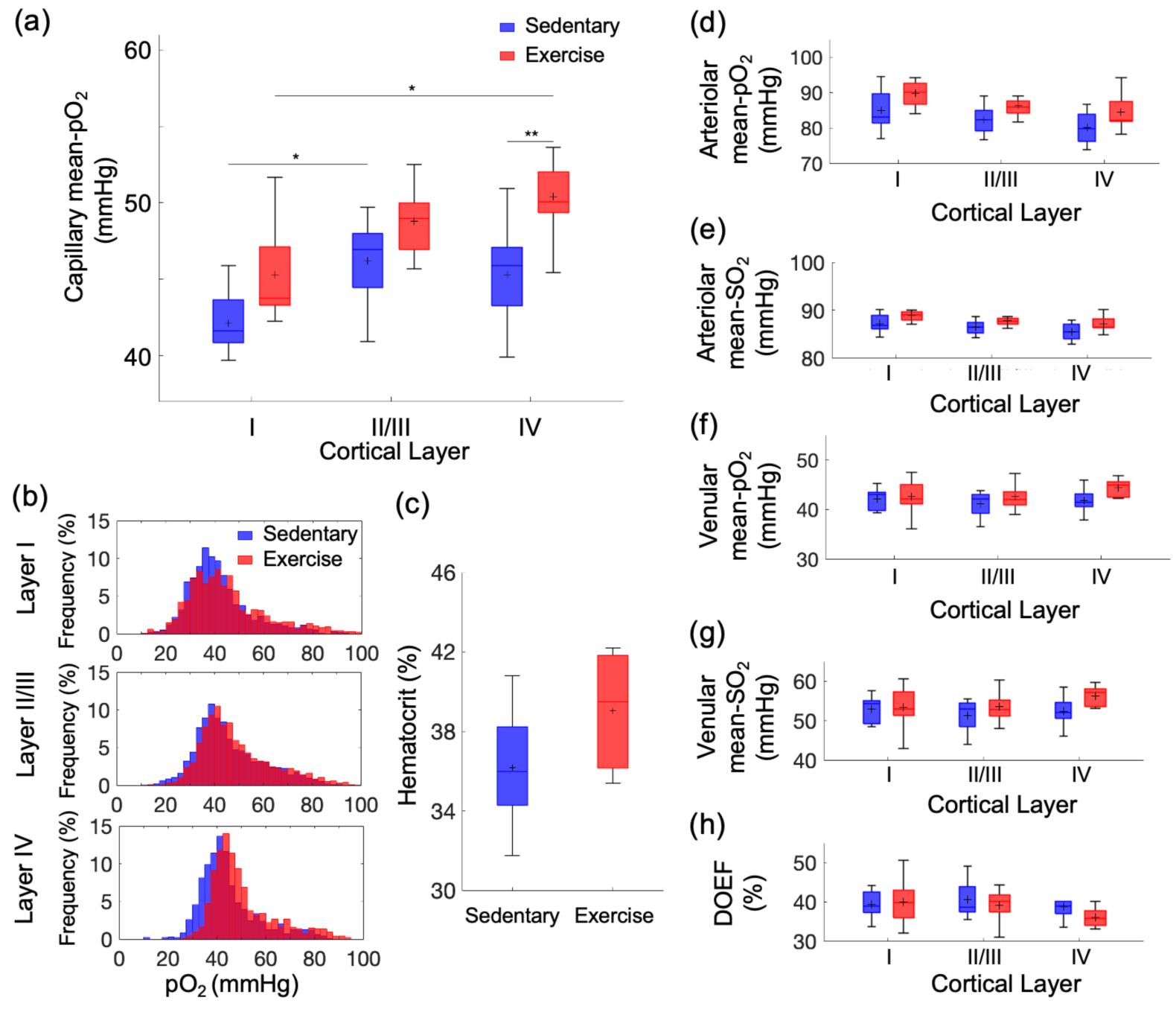
Exercise-induced changes in microvascular pO_2_ across cortical layers in old mice. (a) Capillary mean-pO_2_ across cortical layers in sedentary controls and exercising mice. (b) Histograms of capillary pO_2_ in layers I, II/III, and IV. (c) The mean Hct levels from sedentary (n=8) and exercise group (n=5). (d) and (e) Intravascular pO_2_ and SO_2_ in the diving arterioles across cortical layers I-IV in sedentary (blue boxplots) and exercise (red boxplots) group, respectively. (f) and (g) Intravascular pO_2_ and SO_2_ in the surfacing venules across cortical layers I-IV in sedentary (blue boxplots) and exercise (red boxplots) group, respectively. (h) Depth-dependent OEF in sedentary (blue boxplots) and exercise (red boxplots) group. The analysis in (a) and (b) was made with 1224, 2601, and 922 capillaries across n=9 mice in sedentary group and 1334, 2840, and 1078 capillaries across n=9 mice in exercise group in cortical layers, I, II/III, and IV, respectively. The analysis in (d-h) was made with 13 arterioles and 12 venules from n=9 mice in sedentary group and 14 arterioles and 12 venules from n=9 mice in exercise group. Statistical analysis was carried out using Two-way ANOVA with post hoc Tukey’s in (a) and (d-h) and Student’s t-test in (f). *p<0.05; **p<0.01. Additional details on boxplots and exclusions are provided in the Supplementary document.

The exercise group showed an overall increase in capillary mean-pO_2_ in comparison with the sedentary group across all cortical layers (Figure 4a). The pO_2_ increase in cortical layer IV was more pronounced compared to the other layers. The distribution of capillary pO_2_ in the exercise group also shifted towards higher pO_2_ compared with sedentary controls (Figure 4b), which was again more pronounced in layer IV. Consistent with our previous report,(Li et al., n.d.) we also found a strong positive correlation between the mean pO_2_ and mean capillary RBC flux in both groups (Figure 4 – Figure supplement 1).

The increase in the capillary pO_2_ could be in part due to an increase in Hct level due to exercise.(Moeini et al., 2020) We observed a moderate but statistically insignificant increase in Hct in the exercise group compared with sedentary mice (Figure 4c). No difference in the capillary RBC line-density between two groups was found (Figure 3 – Figure supplement 2). No correlation between the measured parameters (mean pO_2_ and Hct) and the average daily running distance was found (data not shown).

The pO_2_ and SO_2_ in the diving cortical arterioles and ascending venules across cortical layers were lower in the old sedentary mice (Figure 4d-g) than what we have reported in young adult mice,(Li et al., n.d.) and in agreement with the pO_2_ values in old mice previously reported by others.(Moeini et al., 2018) Importantly, both pO_2_ and SO_2_ tended to increase in all cortical layers in exercise group compared with sedentary group, particularly in layer IV, consistent with the largest changes in the capillary RBC flux and mean pO_2_ observed in the deeper cortical layers. Consequently, the depth-dependent oxygen extraction fraction (DOEF) decreased in the exercise group compared with the sedentary controls, with potentially the largest decrease in the layer IV (Figure 4h).

### 3.3 Cortical hemodynamic response to functional activation was reduced by aging but not altered by exercise

The subtle impact of exercise on the gray matter microvascular perfusion and oxygenation led us to question whether it has effects on cortical hemodynamic response. We assessed the effects of aging and exercise on the hemodynamic response to the whisker stimulation using OISI (Figure 5). Optical intrinsic signal time courses of mice in the exercise and sedentary group were obtained (Figure 5a) and compared with those of younger (7 months old) sedentary mice (Figure 5b). The peak amplitudes of the hemodynamic response were significantly smaller in aged mice compared with younger mice regardless of the exercise status (Figure 5c). However, the exercise did not affect the peak response amplitude. Response latency (i.e., time to peak after stimulus onset) also did not differ among the groups (Figure 5 – Figure supplement 1). No correlation between the measured parameters (the peak response amplitude and the response latency) and the average running distance was found (data now shown).

**Figure 5.**
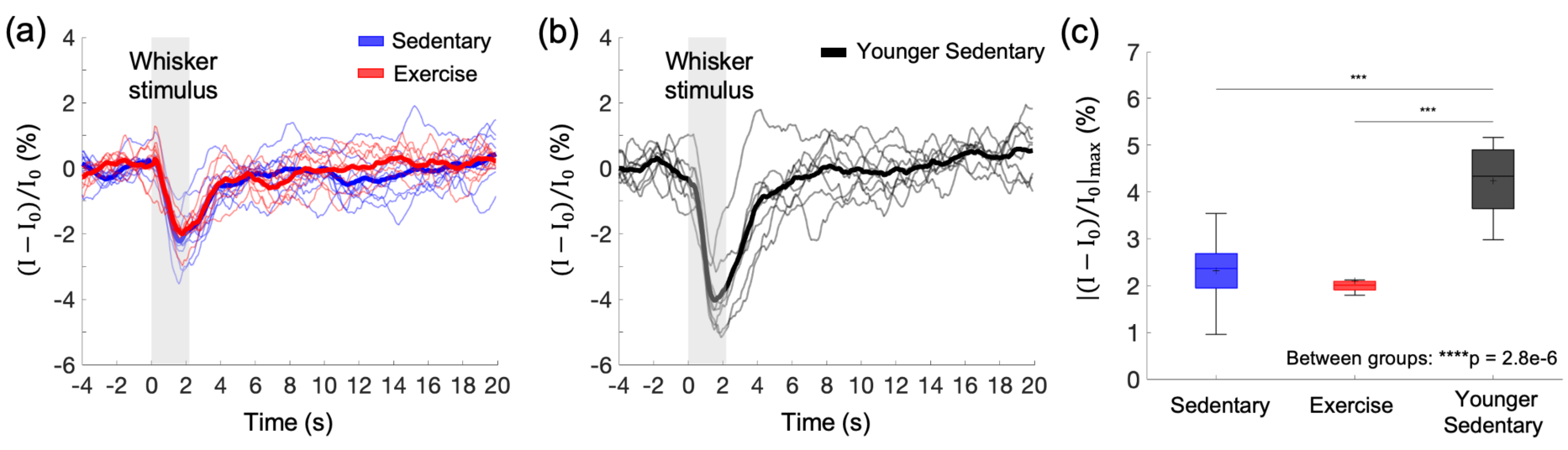
Effects of aging and exercise on functional hemodynamic response. (a, b) Optical intrinsic signal time courses in the whisker barrel cortex of individual old mice in aged sedentary (blue; n=9) and exercise (red; n=8) mice (a), and younger (7 months old) sedentary mice (n=8) (b). Thick curves represent averages. (c) Average changes in the peak intensity in old sedentary, old exercise, and young group. One-way ANOVA with Tukey post hoc test. *** p<0.001. Please see Supplementary document for exclusions.

### 3.4 Cortical microvascular density is significantly larger in the exercise group

To explore whether excise induces structural changes in the cerebral microvasculature of aged mice, we segmented three-dimensional stacks of the cortical microvasculature and obtained their mathematical graph representations.(Tsai et al., 2009) Representative maximum intensity projection (MIP) images of the microvascular stacks show denser cortical microvascular networks of mice from the exercise group compared to those of the sedentary controls (Figure 6a). Exercise group had significantly higher microvascular segment and length density compared with their sedentary controls (Figure 6b, c). Segment and length density did not correlate with running activity (data not shown).

**Figure 6.**
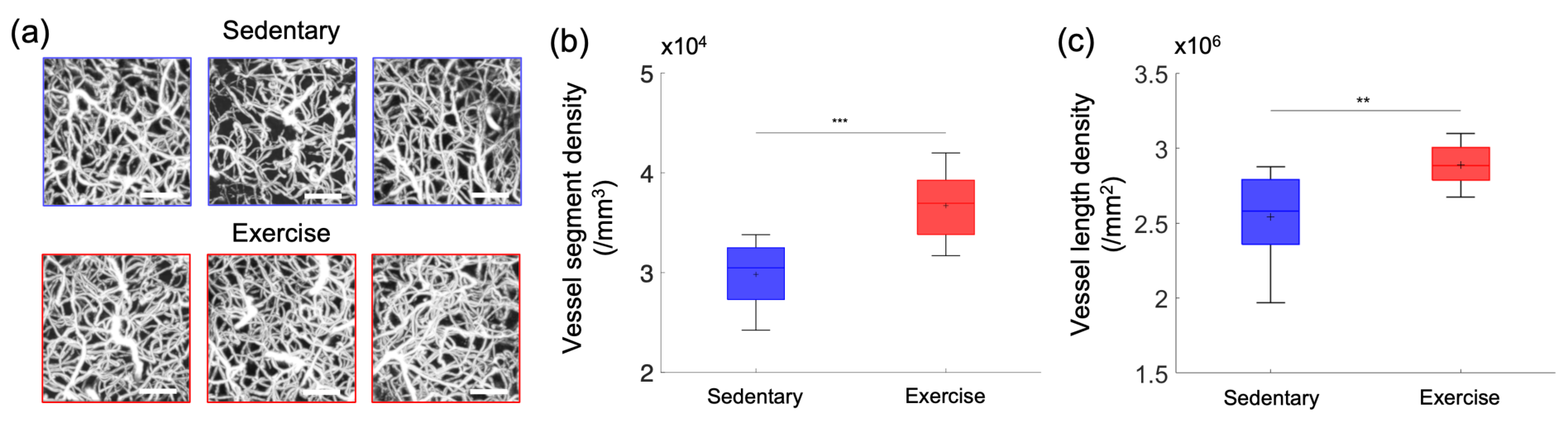
Cortical microvascular density in aged mice. (a) Representative MIP images of the three-dimensional angiograms of three mice in the sedentary group and three mice in the exercise group, over the cortical depth range from 50 µm to 400 µm, and 200 × 200 µm^2^ FOV. Scale bars: 50 µm. (b) Vessel segment density and (c) vessel length density of cortical capillaries from sedentary (n=19,669 segments; n=9 mice) and exercise (n=18,044 segments; n=8 mice) group. Student’s t-test. **p<0.01; *** p<0.001. Please see Supplementary document for exclusions.

### 3.5 Exercise improves short-term spatial memory performance

In the novel object recognition test, mice from the exercise group spent more time exploring the novel object than the familiar one leading to significantly higher DI scores across four different time intervals than sedentary mice (Figure 7a). Interestingly, the DI score for each time interval correlated with the average daily running distance (Figure 7b-e). In contrast, sedentary and exercise groups did not differ in the Y-maze test performance (Figure 7f).

**Figure 7.**
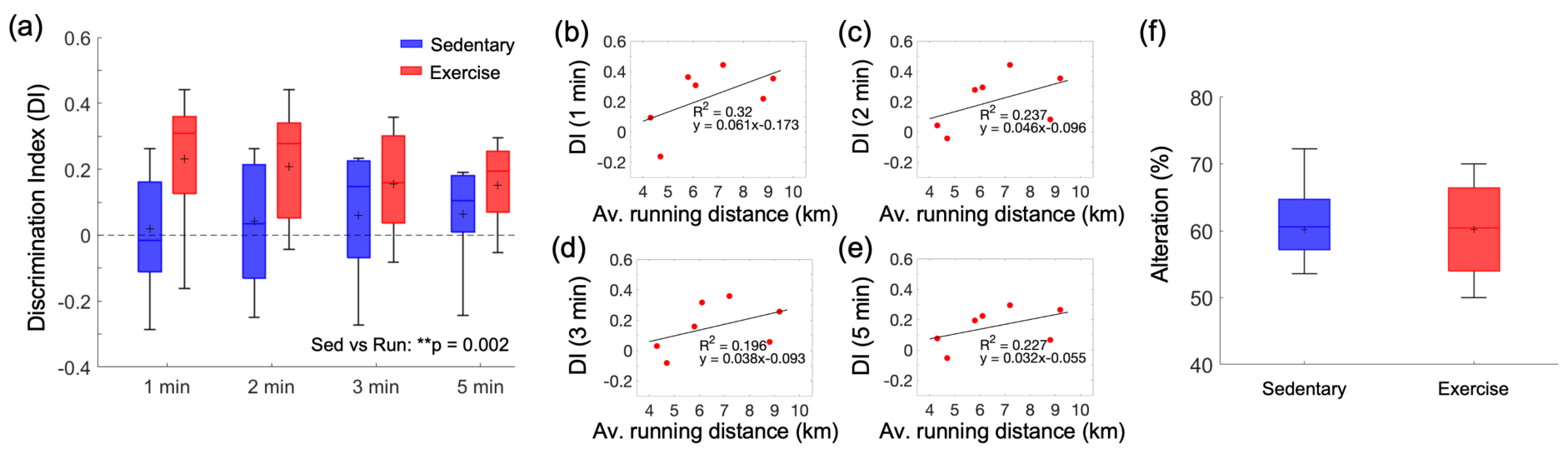
Effect of exercise in the old mice on cognitive performance. (a) DI scores in NORT, calculated with four different exploration time periods in the sedentary (n=9) and exercise (n=7) group. The calculated DI values at each time interval were subsequently averaged across animals. (b-d) Correlations between the daily average running distance and four different DI scores: 1, 2, 3, and 4 minutes, respectively. (f) Spontaneous alteration scores in the Y-maze test in the sedentary (n=9) and exercise (n=7) group. Statistical analysis was carried out using Two-way ANOVA with post hoc Tukey’s in (a) and Student’s t-test in (f). **p<0.01. Please see Supplementary document for exclusions.

## 4. Discussion

In this study, we observed depth-dependent decreases in cerebral microvascular perfusion and oxygenation in aged sedentary mice. The decrease was mitigated by five months of exercise in a depth-dependent manner. The key findings include (1) age-associated reduction in capillary RBC flux in the white matter that was moderated by exercise; (2) a decrease in the mean capillary pO_2_ in cortical layer IV with aging and the overall improvement in capillary pO_2_ with exercise, with a particularly pronounced pO_2_ increase in layer IV; (3) an increase in cortical microvascular density with exercise; and (4) improvement of short-term memory function with exercise.

We first assessed the spatial distribution of capillary RBC flux across the cortical and subcortical regions and how it responds to aging and exercise. In aged sedentary mice, the capillary RBC flux in the subcortical white matter tended to be lower than that in the gray matter (Figure 3b). Young adult sedentary mice showed the opposite trend (Figure 3a), consistent with our previous finding in anesthetized young adult mice (3-5 months old) showing a higher capillary RBC flux in the subcortical white matter compared to the gray matter.(Li et al., 2020) Importantly, we observed a large discrepancy in the capillary RBC flux between young and aged sedentary mice in the gray matter and an almost two-fold greater RBC flux discrepancy between the two groups in the white matter. Cerebral white matter is known to be more susceptible to hypoperfusion compared to the gray matter, potentially because it is located further downstream with respect to the arterial blood supply.(Gunning-Dixon et al., 2009; Markus et al., 2000) White matter vulnerability to ischemic injury increases with age,(Baltan et al., 2008) and white matter lesions and lacunar infarcts are common in elderly people with CSVD and hypertension.(Breteler et al., 1994; van Swieten et al., 1991) The vulnerability of white matter to such pathological conditions could be related to our observation of a severe decrease in blood flow in the white matter during normal aging.

Five months of exercise restored the spatial distribution trend of capillary RBC flux close to its distribution in the young sedentary mice (Figure 3b). Increased capillary perfusion in the white matter was associated with increased blood flow in ascending venules, as measured by Doppler OCT (Figure 3e,f). In contrast to smaller ascending venules having their branches being distributed mostly over the upper cortical layers, large venules likely extend further to subcortical white matter where the increased capillary perfusion was observed.(Duvernoy et al., 1981; Kirst et al., 2020; Xiong et al., 2017) Venules smaller than 20 *μm* in diameter were excluded from analysis as the small venules tend to have slower flow and may have smaller absolute changes in the blood flow compared to larger venules with higher flow and thus the flow change may not be easily distinguishable due to the relatively low precision of the speed measurement based on Doppler OCT (±0.7 mm/s, see *Methods 2.4* for more details).(Fan et al., 2020; Santisakultarm et al., 2012)

Previous studies in mice reported conflicting results regarding the effects of exercise on cortical microcirculation.(Dorr et al., 2017; Falkenhain et al., 2020; Lu et al., 2020) However, these observations have been limited to the blood flow in gray matter, which was shown in this study to be less responsive to exercise than that in deeper brain regions. Human MRI data acquired after short-term (one week) exercise showed that exercise induced a selective increase in hippocampal CBF with no or negligible changes in the gray matter CBF.(Steventon et al., 2021) Our findings suggest that cerebral subcortical microcirculation is more responsive to both age-related and exercise-induced changes than cortical microcirculation.

The change in cortical microvascular oxygenation was also depth-dependent. In aged sedentary mice, the mean capillary pO_2_ increased from layer I to layer II/III and reached a plateau in layer II/III (Figure 5a, b). A different trend was observed in young sedentary mice, which showed a gradual increase in the mean capillary pO_2_ from layer I to layer IV.(Li et al., n.d.) In contrast to aged sedentary controls, mice from the exercise group showed a similar trend to young sedentary mice, with a pronounced increase in the mean capillary pO_2_ in layer IV among all assessed cortical layers. In the mouse somatosensory cortex, layer IV exhibits the highest neuronal and capillary densities and the strongest staining for the cytochrome c oxidase, potentially implying the highest oxidative demand during activation and/or at rest throughout all cortical layers.(Blinder et al., 2013; Lefort et al., 2009) In an immunohistochemical study performed using an anti-Glut-1 antibody, high plaque load and decreased blood vessel density were observed, particularly in layer IV of the somatosensory cortex in aged, 18-month-old transgenic AD mice, while younger AD mice did not demonstrate any difference compared to the wild-type mice.(Kuznetsova & Schliebs, 2013) The decrease in capillary mean pO_2_ in layer IV in aged sedentary mice could be associated with the larger mismatch in the oxygen delivery and consumption compared with the more superficial cortical layers. However, due to technical limitations, we did not assess the microvascular oxygenation in the deeper cortical layers and subcortical white matter. Because RBC flux exhibits the largest discrepancy between sedentary old and young mice in the subcortical white matter, it is possible that intravascular pO_2_ also exhibits the largest discrepancy in this brain region.

Voluntary exercise significantly improved microvascular oxygenation compared to sedentary controls, particularly in layer IV. Similar to our finding of capillary RBC flux improvement due to exercise mostly in the white matter, exercise differentially affected cortical intravascular oxygenation, with the largest increase in the deeper layers and restored the spatial distribution trend of capillary pO_2_ across cortical layers close to its distribution in the young sedentary mice. However, since the difference in both the Hct level and RBC line density between the two groups was not significant (Figure 5c and Figure 3 – Figure supplement 2), factors other than increased Hct may be involved in the observed depth-dependent pO_2_ increase in the gray matter. Interestingly, we found that the RBC line density in both groups of aged mice was significantly higher in the subcortical white matter than in the gray matter. This was not observed in young adult mice.(Li et al., 2020) However, this finding is consistent with previous simulation results that showed higher Hct levels in deep-reaching penetrating arterioles compared to arterioles whose branches connect to the capillary bed closer to the surface due to the plasma skimming effect.(Gould et al., 2017)

We observed no change in the relative peak amplitude or the latency of stimulus-induced hemodynamic response with exercise, assessed with OISI at 570 nm, which emphasizes the intrinsic signal originating from cerebral blood volume changes in the superficial cortical layers (Figure 5 and Figure 5 – Figure supplement 1).(Malonek et al., 1997; Tian et al., 2011) Young sedentary mice (7 months old) showed a significantly larger relative response amplitude than aged mice, consistent with the reduced cerebrovascular reactivity with age in healthy adults and rodents.(Bálint et al., 2019; J. N. Barnes, 2015; Seker et al., 2021) The baseline CBF can have a strong effect on the magnitude of the hemodynamic response.(Buxton et al., 2004; Corfield et al., 2001) While no statistically significant difference in the mean capillary RBC flux in the gray matter was found between the aged sedentary and exercise group (Figure 3b), the exercise group had larger capillary density in the gray matter (Figure 6b), suggesting that cortical blood perfusion was possibly also higher in the exercise group. This was further supported by significantly higher blood flow in ascending venules in the same group (Figures 3e,f). Therefore, our data implies that with a similar level of the relative response amplitude between two groups, the exercise group potentially had a larger transient blood supply to the tissue during functional hyperemia.

Increased capillary density in the sensorimotor cortex and hippocampus in response to exercise has been observed in both old rats and young adult mice.(Ding et al., 2006; Morland et al., 2017) However, other studies showed no improvement in cerebral microvascular structure with regular exercise in the sensorimotor cortex of mice of a similar age range.(Dorr et al., 2017; Falkenhain et al., 2020) Although it is unclear whether the difference in age or brain region (or both) contributed to the inconsistent results with some of the previous observations, we found significant morphological changes in the microvascular morphometric parameters due to chronic exercise in the somatosensory cortex of 20-month-old mice (Figure 6). Based on the observed changes in the microvascular RBC flux and pO_2_, we anticipate that an even greater discrepancy in the capillary density between the two groups of mice may be observed in deeper cortical layers and subcortical white matter. It is compelling to hypothesize that higher microvascular density is one of the major contributors to the larger microvascular perfusion and oxygenation observed in mice from the exercise group. However, it is not clear whether the larger capillary density in the exercise group is mostly due to increased angiogenesis or decreased pruning of the capillaries in comparison with the sedentary group. In addition, if increased angiogenesis is present in the exercise group, it will be important to better understand the contribution of the new capillaries to oxygen delivery to tissue.

Increased cerebral perfusion and oxygenation with exercise were accompanied by improved spatial short-term memory function, as evaluated by NORT (Figure 7a). The perirhinal cortex plays an important role in object recognition memory. It receives sensory inputs from its neighboring sensory cortices, such as the somatosensory cortex, where we found significant improvements in vascular function and structure due to exercise in aged mice.(Antunes & Biala, 2012; Cohen & Stackman Jr., 2015) In contrast, we did not observe an increase in spontaneous alterations between the arms of the Y-maze (Figure 7c). As previously shown in several studies in mouse models of AD, each memory task depends on a variety of brain regions that can be differentially affected by exercise, which could potentially produce inconsistent results between different memory tasks.(Kraeuter et al., 2019; Winters, 2004) The NORT results depend on the exploration time in the test phase, which is related to the age-dependent decay of the novel object preference.(Traschütz et al., 2018) Therefore, choosing an adequate time interval to calculate the discrimination index is important to reliably detect novel object recognition, especially in aged mice. In our analysis, the DI score was calculated at four different time intervals: 1, 2, 3, and 5 min. The results showed an increased DI score in running mice compared to sedentary mice across all time intervals, which was also positively correlated with the average running distance (Figure 7b).

A recent study reported a positive correlation between the average daily running distance and cerebral tissue oxygenation in 6-month-old mice after three months of exercise.(Lu et al., 2020) While mice in our exercise group had significantly higher white matter capillary RBC flux than sedentary mice, the mean capillary RBC flux in the white matter was negatively correlated with the average daily running distance (Figure 3 – Figure supplement 3). Although white matter capillary density was not assessed in this study, lower capillary RBC flux in the white matter could potentially be accompanied with higher capillary density. In this case, capillary RBC flux in the white matter can decrease but cerebral tissue oxygenation in this region can still increase in response to chronic exercise. This result also may be due to a non-proportional relationship between exercise and brain function, which could decrease after reaching the optimal intensity of exercise.(Khakroo Abkenar et al., 2019) It is possible that five months of unrestricted exercise led some (or all) animals to over-exercise, which may adversely affect subcortical capillary blood flow and prevent achieving the maximum benefits of exercise on the microcirculation.

In the analysis of the capillary RBC flux and capillary pO_2_, all capillaries identified within the FOV were selected and used for analysis without considering their branching orders. In future studies, analysis based on different types of vessels classified by branching order could potentially provide additional insight into capillary blood flow and oxygenation, as we previously showed different characteristics of capillaries with low branching orders located close to precapillary arterioles compared with higher-order capillaries.(Li et al., n.d.)

In conclusion, leveraging our multimodal optical imaging tools, we quantified the changes in microvascular function, structure, and sensory-evoked functional hyperemia in response to aging and voluntary exercise in 20-month-old mice. Our results indicate that cerebral microcirculation and oxygenation in deeper cortical layers and subcortical white matter are more susceptible to age-related degeneration, but they are surprisingly more responsive to voluntary aerobic exercise, which induces significant improvements in capillary density, RBC flux, and intracapillary pO_2_ . Improvements in cerebrovascular function and structure are accompanied by rescue of cognitive function. These findings may help us to better understand the patterns and consequences of age-related decline of microcirculation at different cortical depths and subcortical white matter, and how the neurologic effects of aging may be ameliorated by aerobic exercise.

## Acknowledgements

Support of the grants RF1NS121095, R01NS115401, U24EB028941, U19NS123717, U01HL133362, R00MH120053, and R01AG055413 from the National Institutes of Health, USA, and support from the Rappaport Foundation and Leducq Foundation are gratefully acknowledged. The funders had no role in study design, data collection and interpretation, or the decision to submit the work for publication.

## 6. Competing interests

The authors declare no competing interests.

**[Source Files]**

**Figure3_SourceData1**

Capillary RBC flux measured in young sedentary mice

**Figure3_Source Data2**

Capillary RBC flux measured in aged mice

**Figure3_Source Data3**

Venous flow measured in aged mice

**Figure4_Source Data1**

Capillary pO_2_ measured in aged mice

**Figure4_Source Data2**

Blood hematocrit level

**Figure4_Source Data3**

Arterial (and venous) pO_2_ measured in aged mice

**Figure5_Source Data1**

Peak hemodynamic response amplitude measured in aged mice

**Figure6_Source Data1**

Cortical capillary segment/length density measured in aged mice

**Figure7_Source Data1**

Behavioral scores from NORT and Y-maze test

## Supplementary Figures

**Figure 3 – Figure supplement 1.**
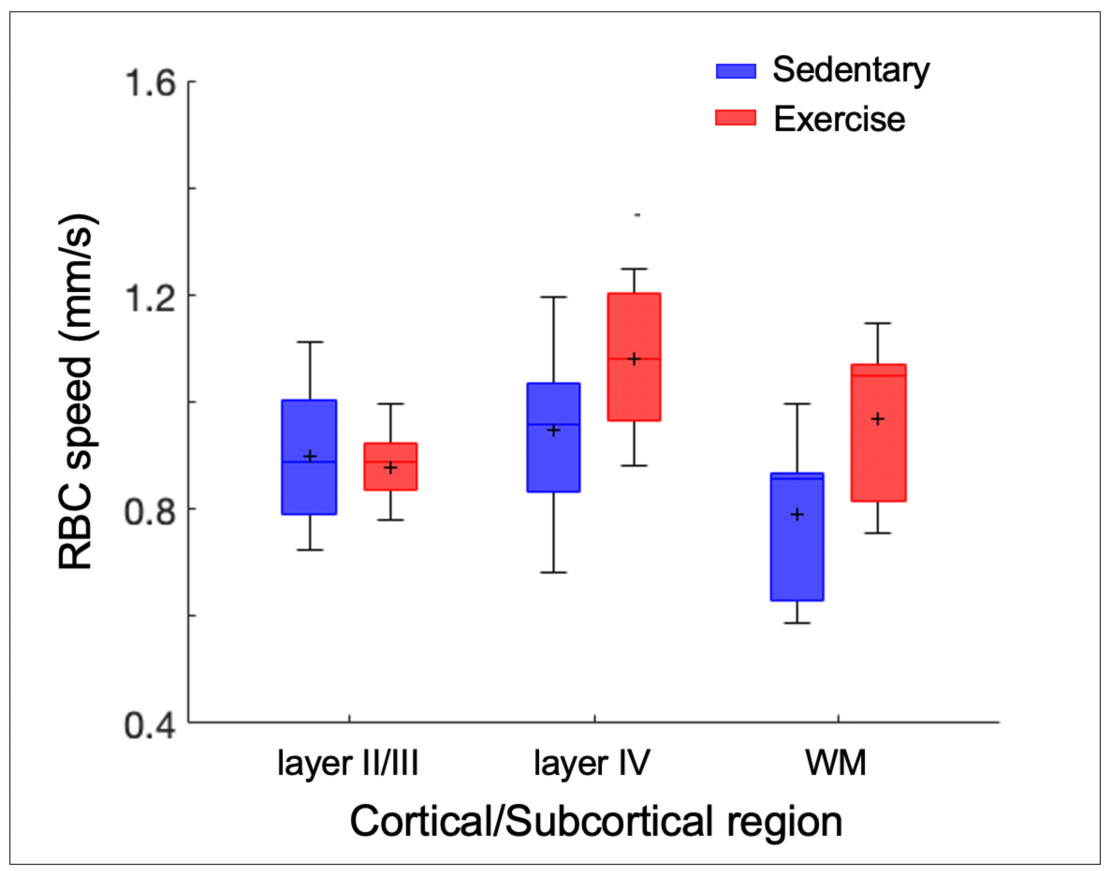
Exercise induced alterations in capillary RBC speed. Capillary RBC speed across cortical layers II/III and IV, and subcortical white matter in aged sedentary and running mice, respectively. The analysis was made with 921, 486, and 112 capillaries across n=7 mice in the sedentary group and 1046, 465, and 238 capillaries across n=8 mice in the exercise group in cortical layers, II/III, IV, and the white matter. Statistical comparisons were carried out using Two-way ANOVA with Tukey post hoc test.

**Figure 3 – Figure supplement 2.**
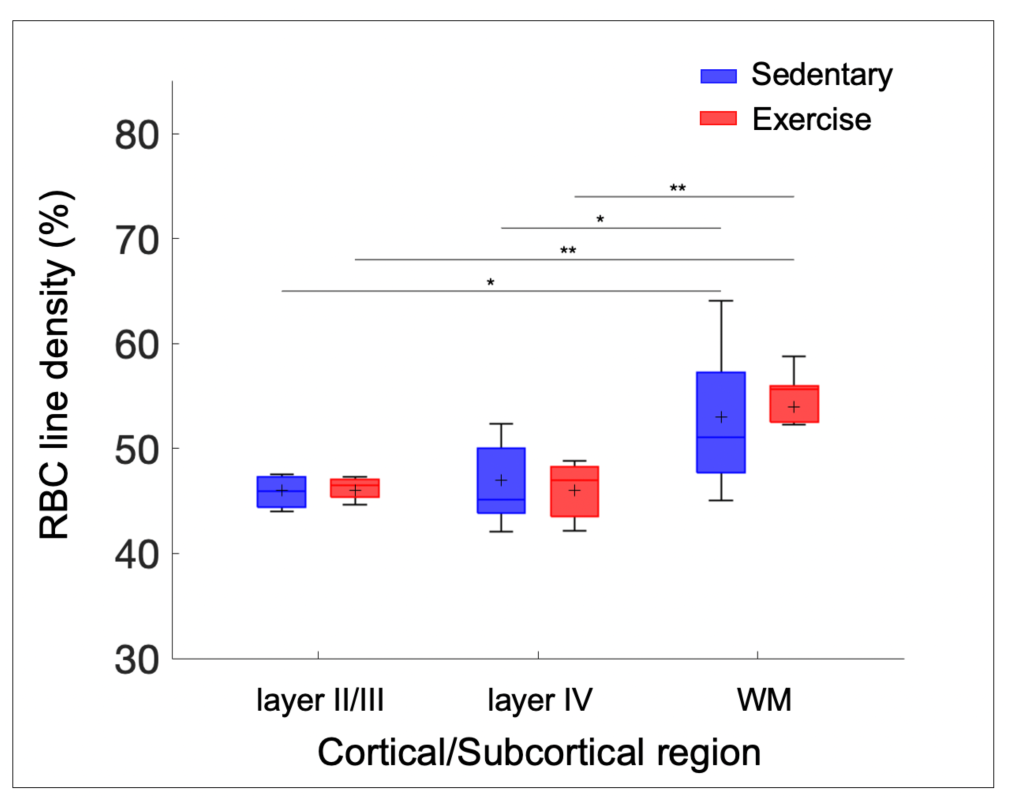
Exercise induced alterations in capillary RBC line-density. Capillary RBC line-density across cortical layers II/III and IV, and subcortical white matter in aged sedentary and running mice. The analysis was made with 921, 486, and 112 capillaries across n=7 mice in sedentary group and 1046, 465, and 238 capillaries across n=8 mice in the exercise group in cortical layers, II/III, IV, and the white matter. Statistical comparisons were carried out using Two-way ANOVA with Tukey post hoc test. The single-asterisk symbol (*) indicates p<0.05; the double-asterisk symbol (**) indicates p<0.01.

**Figure 3 – Figure supplement 3.**
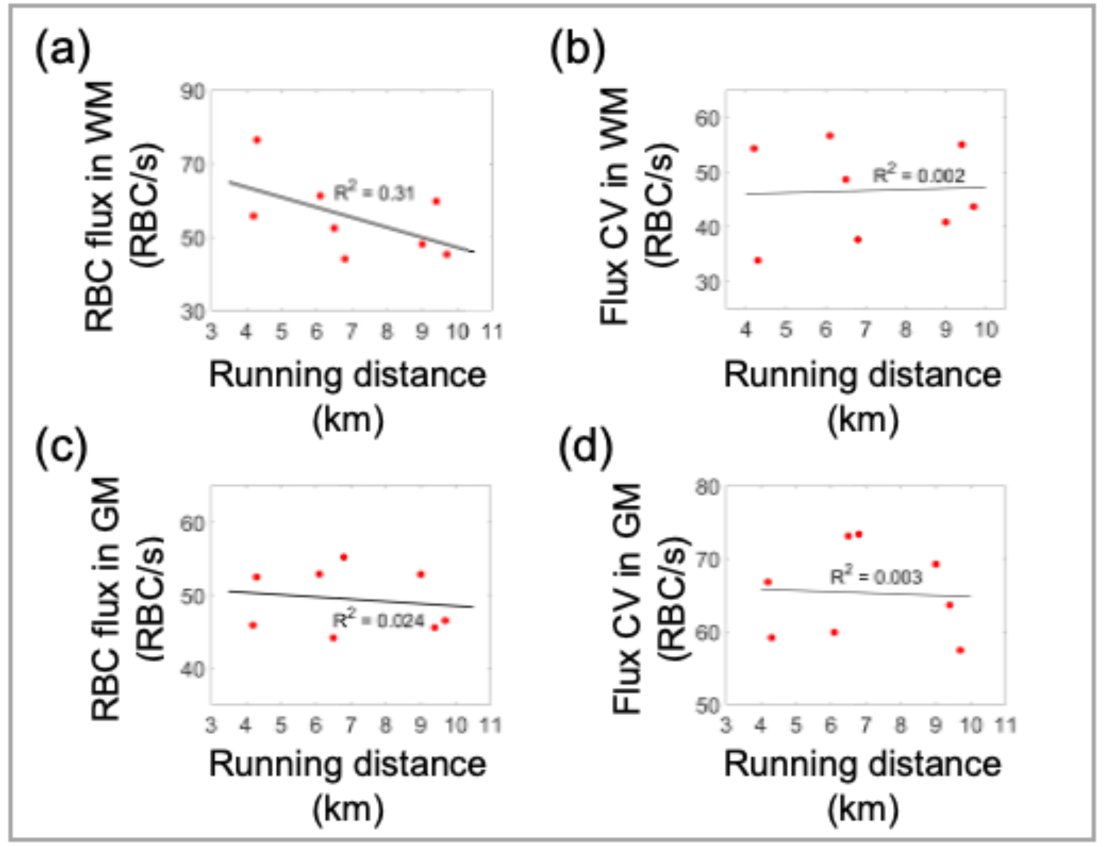
Correlations between the capillary RBC flux and CV, and the running activity. Each data point represents an individual animal. The black solid line is the best linear regression fit. (a) and (b) Correlations between the capillary RBC flux and CV in the white matter and the average daily running distance, respectively. (c) and (d) Correlations between the CV of capillary RBC flux and CV in the gray matter and the average daily running distance, respectively. For each animal, the gray matter capillary RBC flux was calculated by averaging the acquired flux values from cortical layers II/III and IV.

**Figure 4 – Figure supplement 1.**
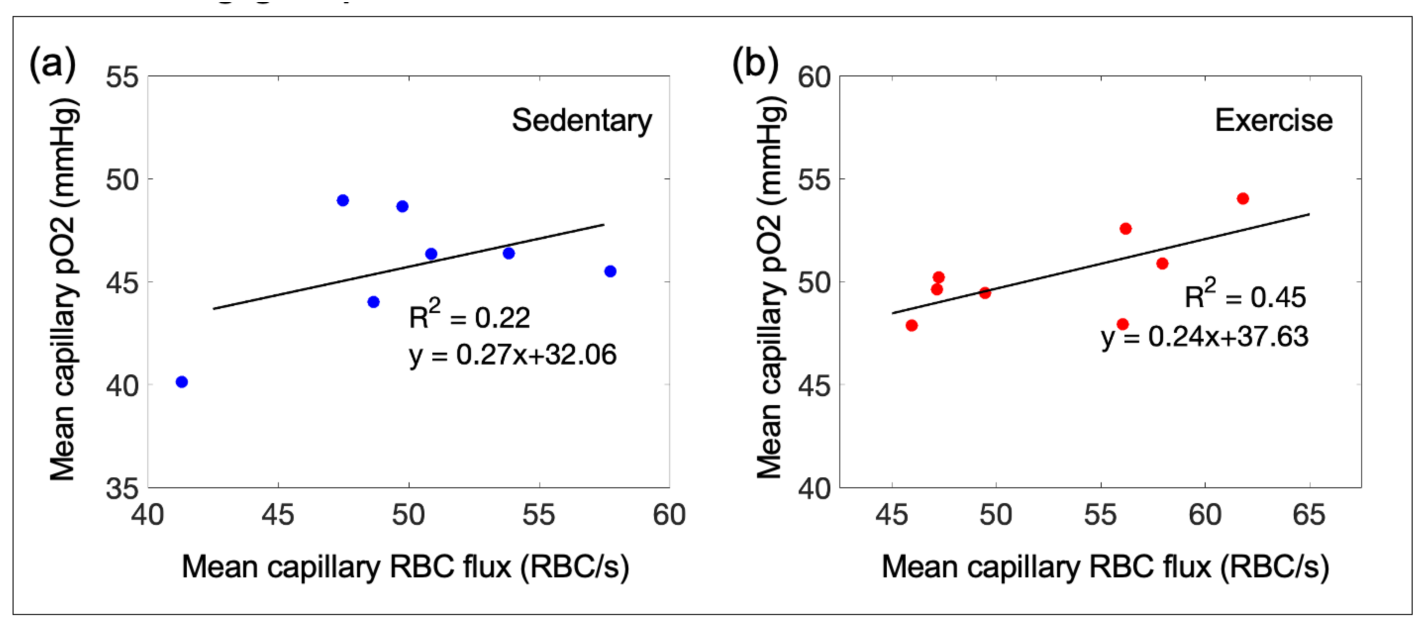
Capillary mean-*pO*_2_ vs capillary RBC flux in the mouse cortex. (a) and (b) Correlations between the mean capillary *pO*_2_ and the mean capillary RBC flux in sedentary and running mice, respectively. The black solid line is the best fit result of each linear regression (R^2^ = 0.22, *y* = 0.27*x* + 32.06, for sedentary mice and R^2^ = 0.45, *y* = 0.24*x* + 37.63, for running mice). Each data point represents an individual animal. For each animal, the mean capillary RBC flux and the mean capillary *pO*_2_ were calculated by averaging the acquired RBC flux and *pO*_2_ values from cortical layers II/III and IV, and cortical layers I, II/III, and IV, respectively. The correlation coefficient (the R value) for each group was converted to Fisher z value to compare difference between correlation coefficients of sedentary and running mice, and no significant difference was found (the observed z value = 0.44).

**Figure 5 – Figure supplement 1.**
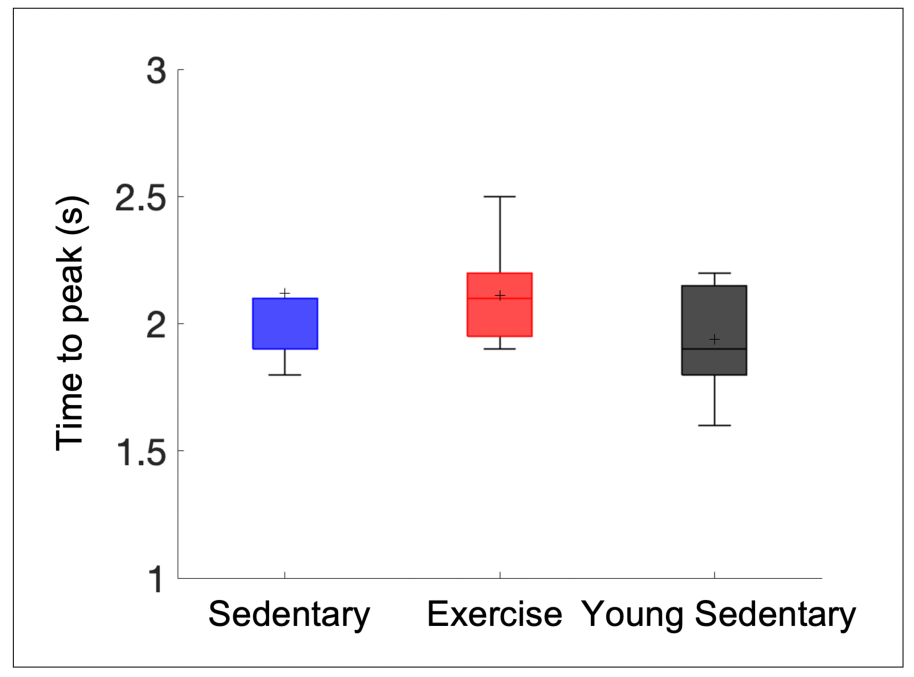
Differences in the latency (time to peak) of stimulus-induced hemodynamic response between sedentary, running, and younger mice. Statistical comparisons were carried out using One-way ANOVA with Tukey post hoc test.

## Reference

Antunes, M., & Biala, G. (2012). The novel object recognition memory: neurobiology, test procedure, and its modifications. Cognitive Processing, 13(2), 93–110. https://doi.org/10.1007/s10339-011-0430-z

Bálint, A. R., Puskás, T., Menyhárt, Á., Kozák, G., Szenti, I., Kónya, Z., Marek, T., Bari, F., & Farkas, E. (2019). Aging Impairs Cerebrovascular Reactivity at Preserved Resting Cerebral Arteriolar Tone and Vascular Density in the Laboratory Rat. Frontiers in Aging Neuroscience, 11. https://doi.org/10.3389/fnagi.2019.00301

Baltan, S., Besancon, E. F., Mbow, B., Ye, Z., Hamner, M. A., & Ransom, B. R. (2008). White Matter Vulnerability to Ischemic Injury Increases with Age Because of Enhanced Excitotoxicity. Journal of Neuroscience, 28(6), 1479–1489. https://doi.org/10.1523/JNEUROSCI.5137-07.2008

Barnes, D. E., Yaffe, K., Satariano, W. A., & Tager, I. B. (2003). A Longitudinal Study of Cardiorespiratory Fitness and Cognitive Function in Healthy Older Adults. JAGS, 51, 459–465.

Barnes, J. N. (2015). Exercise, cognitive function, and aging. Advances in Physiology Education, 39(2), 55–62. https://doi.org/10.1152/advan.00101.2014

Blinder, P., Tsai, P. S., Kaufhold, J. P., Knutsen, P. M., Suhl, H., & Kleinfeld, D. (2013). The cortical angiome: An interconnected vascular network with noncolumnar patterns of blood flow. Nature Neuroscience, 16(7), 889–897. https://doi.org/10.1038/nn.3426

Breteler, M. M. B., van Swieten, J. C., Bots, M. L., Grobbee, D. E., Claus, J. J., van den Hout, J. H. W., van Harskamp, F., Tanghe, H. L. J., de Jong, P. T. V. M., van Gijn, J., & Hofman, A. (1994). Cerebral white matter lesions, vascular risk factors, and cognitive function in a population-based study. Neurology, 44(7), 1246–1246. https://doi.org/10.1212/WNL.44.7.1246

Buxton, R. B., Uludağ, K., Dubowitz, D. J., & Liu, T. T. (2004). Modeling the hemodynamic response to brain activation. NeuroImage, 23, S220–S233. https://doi.org/10.1016/j.neuroimage.2004.07.013

Cees De Groot, J., de Leeuw, F.-E., Oudkerk, M., van Gijn, J., Hofman, A., Jolles, J., & Breteler, M. M. B. (2000). Cerebral White Matter Lesions and Cognitive Function: The Rotterdam Scan Study. Annals of Neurology, 47, 145–157.

Cohen, S. J., & Stackman Jr., R. W. (2015). Assessing rodent hippocampal involvement in the novel object recognition task. A review. Behavioural Brain Research, 285, 105–117. https://doi.org/10.1016/j.bbr.2014.08.002

Corfield, D. R., Murphy, K., Josephs, O., Adams, L., & Turner, R. (2001). Does Hypercapnia-Induced Cerebral Vasodilation Modulate the Hemodynamic Response to Neural Activation? NeuroImage, 13(6), 1207–1211. https://doi.org/10.1006/nimg.2001.0760

de Leeuw, F. E., de Groot, J. C., Achten, E., Oudkerk, M., Ramos, L. M. P., Heijboer, R., Hofman, A., Jolles, J., van Gijn, J., & Breteler, M. M. B. (2001). Prevalence of cerebral white matter lesions in elderly people: A population based magnetic resonance imaging study. The Rotterdam Scan Study. Journal of Neurology Neurosurgery and Psychiatry, 70(1), 9–14. https://doi.org/10.1136/jnnp.70.1.9

de Miguel, Z., Khoury, N., Betley, M. J., Lehallier, B., Willoughby, D., Olsson, N., Yang, A. C., Hahn, O., Lu, N., Vest, R. T., Bonanno, L. N., Yerra, L., Zhang, L., Saw, N. L., Fairchild, J. K., Lee, D., Zhang, H., McAlpine, P. L., Contrepois, K., … Wyss-Coray, T. (2021). Exercise plasma boosts memory and dampens brain inflammation via clusterin. Nature, 600(7889), 494–499. https://doi.org/10.1038/s41586-021-04183-x

Defrancesco, M., Egger, K., Marksteiner, J., Esterhammer, R., Hinterhuber, H., Deisenhammer, E. A., & Schocke, M. (2014). Changes in white matter integrity before conversion from mild cognitive impairment to Alzheimer’s disease. PLoS ONE, 9(8). https://doi.org/10.1371/journal.pone.0106062

Ding, Y.-H., Li, J., Zhou, Y., Rafols, J., Clark, J., & Ding, Y. (2006). Cerebral Angiogenesis and Expression of Angiogenic Factors in Aging Rats after Exercise. Current Neurovascular Research, 3(1), 15–23. https://doi.org/10.2174/156720206775541787

Dorr, A., Thomason, L. A., Koletar, M. M., Joo, I. L., Steinman, J., Cahill, L. S., Sled, J. G., & Stefanovic, B. (2017). Effects of voluntary exercise on structure and function of cortical microvasculature. Journal of Cerebral Blood Flow and Metabolism, 37(3), 1046–1059. https://doi.org/10.1177/0271678X16669514

Duvernoy, H. M., Delon, S., & Vannson, J. L. (1981). Cortical blood vessels of the human brain. Brain Research Bulletin, 7(5), 519–579. https://doi.org/10.1016/0361-9230(81)90007-1

Esipova, T. v., Barrett, M. J. P., Erlebach, E., Masunov, A. E., Weber, B., & Vinogradov, S. A. (2019). Oxyphor 2P: A High-Performance Probe for Deep-Tissue Longitudinal Oxygen Imaging. Cell Metabolism, 29(3),736-744.e7. https://doi.org/10.1016/j.cmet.2018.12.022

Esiri, M. M., Nagy, Z., Smith, M. Z., Barnetson, L., Smith, A. D., & Joachim, C. (1999). Cerebrovascular disease and threshold for dementia in the early stages of Alzheimer’s disease. Lancet, 354(9182), 919–920. https://doi.org/10.1016/S0140-6736(99)02355-7

Falkenhain, K., Ruiz-Uribe, N. E., Haft-Javaherian, M., Ali, M., Catchers, S., Michelucci, P. E., Schaffer, C. B., & Bracko, O. (2020). A pilot study investigating the effects of voluntary exercise on capillary stalling and cerebral blood flow in the APP/PS1 mouse model of Alzheimer’s disease. PLoS ONE, 15(8 August). https://doi.org/10.1371/journal.pone.0235691

Fan, J. L., Rivera, J. A., Sun, W., Peterson, J., Haeberle, H., Rubin, S., & Ji, N. (2020). High-speed volumetric two-photon fluorescence imaging of neurovascular dynamics. Nature Communications, 11(1), 6020. https://doi.org/10.1038/s41467-020-19851-1

Gould, I. G., Tsai, P., Kleinfeld, D., & Linninger, A. (2017). The capillary bed offers the largest hemodynamic resistance to the cortical blood supply. Journal of Cerebral Blood Flow & Metabolism, 37(1), 52–68. https://doi.org/10.1177/0271678X16671146

Guizar-Sicairos, M., Thurman, S. T., & Fienup, J. R. (2008). Efficient subpixel image registration algorithms. In OPTICS LETTERS (Vol. 100, Issue 2).

Gunning-Dixon, F. M., Brickman, A. M., Cheng, J. C., & Alexopoulos, G. S. (2009). Aging of cerebral white matter: A review of MRI findings. In International Journal of Geriatric Psychiatry (Vol. 24, Issue 2, pp. 109–117). John Wiley and Sons Ltd. https://doi.org/10.1002/gps.2087

Hase, Y., Ding, R., Harrison, G., Hawthorne, E., King, A., Gettings, S., Platten, C., Stevenson, W., Craggs, L. J. L., & Kalaria, R. N. (2019). White matter capillaries in vascular and neurodegenerative dementias. Acta Neuropathologica Communications, 7(1), 16. https://doi.org/10.1186/s40478-019-0666-x

Islam, M. R., Valaris, S., Young, M. F., Haley, E. B., Luo, R., Bond, S. F., Mazuera, S., Kitchen, R. R., Caldarone, B. J., Bettio, L. E. B., Christie, B. R., Schmider, A. B., Soberman, R. J., Besnard, A., Jedrychowski, M. P., Kim, H., Tu, H., Kim, E., Choi, S. H., … Wrann, C. D. (2021). Exercise hormone irisin is a critical regulator of cognitive function. Nature Metabolism, 3(8), 1058–1070. https://doi.org/10.1038/s42255-021-00438-z

Khakroo Abkenar, I., Rahmani-nia, F., & Lombardi, G. (2019). The Effects of Acute and Chronic Aerobic Activit on the Signaling Pathway of the Inflammasome NLRP3 Complex in Young Men. Medicina, 55(4), 105. https://doi.org/10.3390/medicina55040105

Kirst, C., Skriabine, S., Vieites-Prado, A., Topilko, T., Bertin, P., Gerschenfeld, G., Verny, F., Topilko, P., Michalski, N., Tessier-Lavigne, M., & Renier, N. (2020). Mapping the Fine-Scale Organization and Plasticity of the Brain Vasculature. Cell, 180(4), 780-795.e25. https://doi.org/10.1016/j.cell.2020.01.028

Koch, E., Walther, J., & Cuevas, M. (2009). Limits of Fourier domain Doppler-OCT at high velocities. Sensors and Actuators, A: Physical, 156(1), 8–13. https://doi.org/10.1016/j.sna.2009.01.022

Kraeuter, A.-K., Guest, P. C., & Sarnyai, Z. (2019). The Y-Maze for Assessment of Spatial Working and Reference Memory in Mice (pp. 105–111). https://doi.org/10.1007/978-1-4939-8994-2_10

Kuznetsova, E., & Schliebs, R. (2013). β-Amyloid, Cholinergic Transmission, and Cerebrovascular System - A Developmental Study in a Mouse Model of Alzheimer’s Disease. Current Pharmaceutical Design, 19(38), 6749–6765. https://doi.org/10.2174/13816128113199990711

Lefort, S., Tomm, C., Floyd Sarria, J. C., & Petersen, C. C. H. (2009). The Excitatory Neuronal Network of the C2 Barrel Column in Mouse Primary Somatosensory Cortex. Neuron, 61(2), 301–316. https://doi.org/10.1016/j.neuron.2008.12.020

Li, B., Esipova, T. v, Sencan, I., Kılıç, K., Fu, B., Desjardins, M., Moeini, M., Kura, S., Yaseen, M. A., Lesage, F., Østergaard, L., Devor, A., Boas, D. A., Vinogradov, S. A., & Sakadž, S. (n.d.). More homogeneous capillary flow and oxygenation in deeper cortical layers correlate with increased oxygen extraction. https://doi.org/10.7554/eLife.42299.001

Li, B., Ohtomo, R., Thunemann, M., Adams, S. R., Yang, J., Fu, B., Yaseen, M. A., Ran, C., Polimeni, J. R., Boas, A., Devor, A., Lo, E. H., Arai, K., & Sakadžić, S. (2020). Two-photon microscopic imaging of capillary red blood cell flux in mouse brain reveals vulnerability of cerebral white matter to hypoperfusion. Journal of Cerebral Blood Flow and Metabolism, 40(3), 501–512. https://doi.org/10.1177/0271678X19831016

Lu, X., Moeini, M., Li, B., de Montgolfier, O., Lu, Y., Bélanger, S., Thorin, É., & Lesage, F. (2020). Voluntary exercise increases brain tissue oxygenation and spatially homogenizes oxygen delivery in a mouse model of Alzheimer’s disease. Neurobiology of Aging, 88, 11–23. https://doi.org/10.1016/j.neurobiolaging.2019.11.015

Malonek, D., Dirnagl, U., Lindauer, U., Yamada, K., Kanno, I., & Grinvald, A. (1997). Vascular imprints of neuronal activity: Relationships between the dynamics of cortical blood flow, oxygenation, and volume changes following sensory stimulation. Proceedings of the National Academy of Sciences, 94(26), 14826– 14831. https://doi.org/10.1073/pnas.94.26.14826

Markus, H. S., Lythgoe, D. J., Ostegaard, L., O’sullivan, M., & Williams, C. R. (2000). Reduced cerebral blood flow in white matter in ischaemic leukoaraiosis demonstrated using quantitative exogenous contrast based perfusion MRI. In J Neurol Neurosurg Psychiatry (Vol. 69).

Moeini, M., Cloutier-Tremblay, C., Lu, X., Kakkar, A., & Lesage, F. (2020). Cerebral tissue pO_2_ response to treadmill exercise in awake mice. Scientific Reports, 10(1), 13358. https://doi.org/10.1038/s41598-020-70413-3

Moeini, M., Lu, X., Avti, P. K., Damseh, R., Bélanger, S., Picard, F., Boas, D., Kakkar, A., & Lesage, F. (2018). Compromised microvascular oxygen delivery increases brain tissue vulnerability with age. Scientific Reports, 8(1). https://doi.org/10.1038/s41598-018-26543-w

Morland, C., Andersson, K. A., Haugen, Ø. P., Hadzic, A., Kleppa, L., Gille, A., Rinholm, J. E., Palibrk, V., Diget, H., Kennedy, L. H., Stølen, T., Hennestad, E., Moldestad, O., Cai, Y., Puchades, M., Offermanns, S., Vervaeke, K., Bjørås, M., Wisløff, U., … Bergersen, L. H. (2017). Exercise induces cerebral VEGF and angiogenesis via the lactate receptor HCAR1. Nature Communications, 8(1), 15557. https://doi.org/10.1038/ncomms15557

Pantoni, L. (2010). Cerebral small vessel disease: from pathogenesis and clinical characteristics to therapeutic challenges. In The Lancet Neurology (Vol. 9, Issue 7, pp. 689–701). https://doi.org/10.1016/S1474-4422(10)70104-6

Radanovic, M., Pereira, F. R. S., Stella, F., Aprahamian, I., Ferreira, L. K., Forlenza, O. V., & Busatto, G. F. (2013). White matter abnormalities associated with Alzheimer’s disease and mild cognitive impairment: A critical review of MRI studies. In Expert Review of Neurotherapeutics (Vol. 13, Issue 5, pp. 483–493). https://doi.org/10.1586/ern.13.45

Reber, J., Hwang, K., Bowren, M., Bruss, J., Mukherjee, P., Tranel, D., & Boes, A. D. (2021). Cognitive impairment after focal brain lesions is better predicted by damage to structural than functional network hubs. Proceedings of the National Academy of Sciences, 118(19). https://doi.org/10.1073/pnas.2018784118/-/DCSupplemental

Sakadžić, S., Roussakis, E., Yaseen, M. A., Mandeville, E. T., Srinivasan, V. J., Arai, K., Ruvinskaya, S., Devor, A., Lo, E. H., Vinogradov, S. A., & Boas, D. A. (2010). Two-photon high-resolution measurement of partial pressure of oxygen in cerebral vasculature and tissue. Nature Methods, 7(9), 755–759. https://doi.org/10.1038/nmeth.1490

Santisakultarm, T. P., Cornelius, N. R., Nishimura, N., Schafer, A. I., Silver, R. T., Doerschuk, P. C., Olbricht, W. L., & Schaffer, C. B. (2012). In vivo two-photon excited fluorescence microscopy reveals cardiac-and respiration-dependent pulsatile blood flow in cortical blood vessels in mice. American Journal of Physiology-Heart and Circulatory Physiology, 302(7), H1367–H1377. https://doi.org/10.1152/ajpheart.00417.2011

Seker, F. B., Fan, Z., Gesierich, B., Gaubert, M., Sienel, R. I., & Plesnila, N. (2021). Neurovascular Reactivity in the Aging Mouse Brain Assessed by Laser Speckle Contrast Imaging and 2-Photon Microscopy: Quantification by an Investigator-Independent Analysis Tool. Frontiers in Neurology, 12. https://doi.org/10.3389/fneur.2021.745770

Şencan, İ., Esipova, T., Kılıç, K., Li, B., Desjardins, M., Yaseen, M. A., Wang, H., Porter, J. E., Kura, S., Fu, B., Secomb, T. W., Boas, D. A., Vinogradov, S. A., Devor, A., & Sakadžić, S. (2022). Optical measurement of microvascular oxygenation and blood flow responses in awake mouse cortex during functional activation. Journal of Cerebral Blood Flow and Metabolism, 42(3), 510–525. https://doi.org/10.1177/0271678X20928011

Smith, E. E., & Markus, H. S. (2020). New Treatment Approaches to Modify the Course of Cerebral Small Vessel Diseases. Stroke, 51(1), 38–46. https://doi.org/10.1161/STROKEAHA.119.024150

Snowdon, D. A., Greiner, L. H., Mortimer, J. A., Riley, K. P., Greiner, P. A., Markesbery, W. R., & Reprints, C. (2011). Brain Infarction and the Clinical Expression of Alzheimer Disease The Nun Study From the Sanders-Brown Center on Aging (Drs. JAMA.

Srinivasan, V. J., Atochin, D. N., Radhakrishnan, H., Jiang, J. Y., Ruvinskaya, S., Wu, W., Barry, S., Cable, A. E., Ayata, C., Huang, P. L., & Boas, D. A. (2011). Optical coherence tomography for the quantitative study of cerebrovascular physiology. Journal of Cerebral Blood Flow and Metabolism, 31(6), 1339–1345. https://doi.org/10.1038/jcbfm.2011.19

Steventon, J. J., Chandler, H. L., Foster, C., Dingsdale, H., Germuska, M., Massey, T., Parker, G., Wise, R. G., & Murphy, K. (2021). Changes in white matter microstructure and MRI-derived cerebral blood flow after 1-week of exercise training. Scientific Reports, 11(1). https://doi.org/10.1038/s41598-021-01630-7

Tian, P., Devor, A., Sakadžić, S., Dale, A. M., & Boas, D. A. (2011). Monte Carlo simulation of the spatial resolution and depth sensitivity of two-dimensional optical imaging of the brain. Journal of Biomedical Optics, 16(1), 016006. https://doi.org/10.1117/1.3533263

Traschütz, A., Kummer, M. P., Schwartz, S., & Heneka, M. T. (2018). Variability and temporal dynamics of novel object recognition in aging male C57BL/6 mice. Behavioural Processes, 157, 711–716. https://doi.org/10.1016/j.beproc.2017.11.009

Tsai, P. S., Kaufhold, J. P., Blinder, P., Friedman, B., Drew, P. J., Karten, H. J., Lyden, P. D., & Kleinfeld, D. (2009). Correlations of Neuronal and Microvascular Densities in Murine Cortex Revealed by Direct Counting and Colocalization of Nuclei and Vessels. Journal of Neuroscience, 29(46), 14553–14570. https://doi.org/10.1523/JNEUROSCI.3287-09.2009

Valenzuela, P. L., Castillo-García, A., Morales, J. S., de la Villa, P., Hampel, H., Emanuele, E., Lista, S., & Lucia, A. (2020). Exercise benefits on Alzheimer’s disease: State-of-the-science. In Ageing Research Reviews (Vol. 62). Elsevier Ireland Ltd. https://doi.org/10.1016/j.arr.2020.101108

van Swieten, J. C., Geyskes, G. G., Derix, M. M. A., Peeck, B. M., Ramos, L. M. P., van Latum, J. C., & van Gijn, J. (1991). Hypertension in the elderly is associated with white matter lesions and cognitive decline. Annals of Neurology, 30(6), 825–830. https://doi.org/10.1002/ana.410300612

Wang, R., & Holsinger, R. M. D. (2018). Exercise-induced brain-derived neurotrophic factor expression: Therapeutic implications for Alzheimer’s dementia. In Ageing Research Reviews (Vol. 48, pp. 109–121). Elsevier Ireland Ltd. https://doi.org/10.1016/j.arr.2018.10.002

Winters, B. D. (2004). Double Dissociation between the Effects of Peri-Postrhinal Cortex and Hippocampal Lesions on Tests of Object Recognition and Spatial Memory: Heterogeneity of Function within the Temporal Lobe. Journal of Neuroscience, 24(26), 5901–5908. https://doi.org/10.1523/JNEUROSCI.1346-04.2004

Xiong, B., Li, A., Lou, Y., Chen, S., Long, B., Peng, J., Yang, Z., Xu, T., Yang, X., Li, X., Jiang, T., Luo, Q., & Gong, H. (2017). Precise Cerebral Vascular Atlas in Stereotaxic Coordinates of Whole Mouse Brain. Frontiers in Neuroanatomy, 11. https://doi.org/10.3389/fnana.2017.00128

Yaffe, K., Barnes, D., Nevitt, M., Lui, L.-Y., & Covinsky, K. (2001). A Prospective Study of Physical Activity and Cognitive Decline in Elderly Women. JAMA Internal Medicine, 161(14), 1703–1708. https://jamanetwork.com/

